# Rosetta Energy Analysis of AlphaFold2 models: Point Mutations and Conformational Ensembles

**DOI:** 10.1101/2023.09.05.556364

**Authors:** Richard A. Stein, Hassane S. Mchaourab

## Abstract

There has been an explosive growth in the applications of AlphaFold2, and other structure prediction platforms, to accurately predict protein structures from a multiple sequence alignment (MSA) for downstream structural analysis. However, two outstanding questions persist in the field regarding the robustness of AlphaFold2 predictions of the consequences of point mutations and the completeness of its prediction of protein conformational ensembles. We combined our previously developed method SPEACH_AF with model relaxation and energetic analysis with Rosetta to address these questions. SPEACH_AF introduces residue substitutions across the MSA and not just within the input sequence. With respect to conformational ensembles, we combined SPEACH_AF and a new MSA subsampling method, AF_cluster, and for a benchmarked set of proteins, we found that the energetics of the conformational ensembles generated by AlphaFold2 correspond to those of experimental structures and explored by standard molecular dynamic methods. With respect to point mutations, we compared the structural and energetic consequences of having the mutation(s) in the input sequence versus in the whole MSA (SPEACH_AF). Both methods yielded models different from the wild-type sequence, with more robust changes when the mutation(s) were in the whole MSA. While our findings demonstrate the robustness of AlphaFold2 in analyzing point mutations and exploring conformational ensembles, they highlight the need for multi parameter structural and energetic analyses of these models to generate experimentally testable hypotheses.

## INTRODUCTION

The results from CASP 14, the competition for de novo structure prediction, revolutionized the field of structural biology with models produced by AlphaFold2 at experimental accuracy.^1^ The subsequent publication of the methodology^2^ and release of prediction databases and pipelines intensified the interest in AlphaFold2 and its use in structural biology. The DeepMind team, in collaboration with EMBL-EBI, released putative structures of the human proteome^3^ along with over 45 proteomes of other species. These initial efforts centered around the generation of protein models adopting a single conformation, epitomized by the presentation of a single model per sequence in the AlphaFold2 database. The subsequent ability to run the AlphaFold2 pipeline using a convenient interface, ColabFold,^4^ expanded the ability to understand this tool and to gauge its application to address a variety of protein structure/function questions.

One application of interest is the generation and characterization of protein conformational ensembles. The initial perception was that AlphaFold2 only generates a single conformation even though there are five independently trained modelers. We have shown that for some proteins it is possible to generate multiple conformations via multiple runs with different random seeds.^5^ We further tailored the pipeline by modifying the MSA by either making changes at specific residues or windows of residues across the protein sequence. These *in silico* mutations were introduced in the whole MSA since we found that modifications of only the input sequence did not yield robust results. The methodology has been applied by others to the Major Facilitator Superfamily (MFS) membrane transporters to generate both inward- and outward-facing models.^6^ Other methodologies have also been successful in generating alternate conformations with AlphaFold2, for example using structural templates from homologous sequences^7,8^ or subsampling the MSA.^9^ In particular, MSA subsampling modifies two input parameters, max_msa_clusters and max_extra_msa, to randomly select small subsets of the larger MSA.^9^ A recent preprint has further developed subsampling by clustering the sequences in the MSA leading to multiple smaller input MSAs.^10^ Unlike the unmodified AlphaFold2 algorithm,^11^ application of this subsampling approach promotes conformational sampling of fold-switch proteins. Despite these methodological developments, there is still a perception that AlphaFold2 can only generate a single biologically relevant conformation.^12,13^

Another persistent major question is whether AlphaFold2 can predict the effects of single point mutations. An early short letter concluded that AlphaFold2 fails to predict the effect of destabilizing point mutations. The conclusion was based on the results from three proteins where the AlphaFold2 models of point mutants were folded and similar to the wild-type models, whereas experimentally the mutations induced unfolding.^14^ Other studies have generated AlphaFold2 models of wild-type proteins to extract sequence and physical properties in order to create predictors of mutational effects that do appear to have predictive value.^15–17^ A recently published report found that the internal AlphaFold2 quality metric, pLDDT, was not an adequate measure for the energetic changes due to point mutations.^18^ Conversely, another study found a correlation between pLDDT and pathogenicity in a set of cancer proteins.^19^ In addition, a neural network was able to learn the pathogenicity of BRCA1 mutations from an AlphaFold2 model.^20^ Unfortunately this network was BRCA1 specific, limiting the methodology to targets with already characterized experimental data. While these studies are promising, the dogma remains that AlphaFold2 is not able to predict the effect of point mutations.^21–24^

Here, we combine AlphaFold2 methodologies for modeling point mutations and conformational ensembles with energetic analysis using Rosetta in order to provide a pipeline for model analysis. We expand on our previous work on the exploration of protein conformational space by comparing models generated by AF_cluster to those generated by SPEACH_AF and then relaxing and scoring the energetics of the models with Rosetta. We show that these models map out energy landscapes that track with previously reported conformations and energetics. We extend our observation that a more robust change in the models is obtained if the mutations are placed in the whole MSA compared to just the input sequence. In contrast to previous reports, we find that AlphaFold2 can predict structural changes due to point mutations. However, we demonstrate that an integrated structural/energetic parameterization is necessary to assess the models in the context of the established principle of protein response to mutations. Combined, the results support the fact that AlphaFold2 can generate alternate conformations that span the energy landscape and can predict structural and/or energetic changes due to point mutations.

## METHODS

### Point Mutations

For the BRCT domain of BRCA1, the ubiquitin-associated domain (UBA) of Rad23 (HR23A), the MyUb domain of Myosin VI, isocyanide hydratase, and XylE an MSA was generated with MMSeqs2,^25^ via ColabFold.^4^ The MSA was modified in two ways: the mutation was placed in the input sequence only or across the MSA replacing the amino acid in the same position in the sequence while ignoring gaps.^5^ The parent MSA and the two modified MSAs were input ten times to generate 50 models for each MSA on a locally installed version of ColabFold.^4^ All other parameters were left as default. These models were then compared to either the same domain of the protein from the AlphaFold2 structure database or the appropriate experimental structures using RMSD values or TM-Align.^26^ These models were also minimized and scored with Rosetta using FastRelax with the backbone constrained (Table S1).^27–29^ Outlier Rosetta Energy Scores (RES) were removed if the values exceeded 4 times the standard deviation of the mean. A two-tailed t-test and non-parametric Mann-Whitney U test were carried out with SciPy (v. 1.10.1) to determine whether the TM-scores, RES, or RMSD values for each set of mutant models are different from the set of wild-type models.

To examine a larger data set of mutations, we culled data for bacteriophage T4 Lysozyme (T4L) and the BRCT domain of BRCA1. We chose a variety of substitutions in T4L including all replacements at a single site (Table S2).^30–37^ For BRCA1 we chose 50 mutations within the BRCT domain from the ProteinGym dataset (Table S3).^38,39^ We created 50 fitness bin values that spanned from the highest to lowest fitness values and picked the mutations that were closest to the bin values. As above, MSAs were generated with MMSeqs2,^25^ via ColabFold^4^ and 50 models were generated for each mutation. The effect of the mutations was only examined with the mutation placed across the MSA. These models were scored based on pLDDT values for the whole protein and at the site of mutation, the TM-score and RMSD value to an experimental structure (pdb: 2lzm) for T4 lysozyme and the AlphaFold database structure for BRCA1. For the pLDDT score for the single site, the value was normalized to the average pLDDT value for the residue in the wild-type models. These models were also relaxed and scored with Rosetta as above. To assess whether the different metrics have a monotonic relationship to the ΔΔG of unfolding for T4 lysozyme and to the fitness for the BRCT domain of BRCA1 the Spearman’s rank correlation, ρ, was measured. The magnitude of |ρ| allows for a qualitative description of the correlation between the two parameters.

### Conformational Sampling

The MSAs used were generated with MMSeqs2,^25^ via ColabFold.^4^ The models generated with SPEACH_AF (github.com/RSvan/SPEACH_AF) were those reported previously.^5^ For MSA subsampling, AF_cluster (github.com/HWaymentSteele/AF_Cluster)^10^ was used with the minimum number of sequences per cluster (min_samples) set to 3, 7, or 11. The subsampled MSAs were then used to generate 5 models each using a locally installed version of ColabFold.^4^ The resultant models from both methods were processed to exclude models with a pLDDT less than 70, where a pLDDT of 70 or greater generally corresponds to a correct backbone prediction.^3^ This parsed set of models was further evaluated with principal component analysis (PCA) using ProDy^40^ to remove models that have a high pLDDT but are misfolded or misthreaded. The remaining models were then subjected to minimization in Rosetta utilizing FastRelax with the backbone constrained (Table S1).^27–29^ For the SPEACH_AF models, the residues mutated to alanine were mutated back to the native residues prior to relaxation. The default Rosetta score function was used for soluble proteins and the membrane_highres_Menv_smooth weights were used for membrane proteins. The membrane spanning regions were determined by TOPCONS.^41^ The Rosetta relaxed structures were analyzed with the eigenvalues and eigenvectors from the PCA run on the models prior to relaxation. The Rosetta Energy Scores were adjusted by setting the lowest scoring model to zero, yielding a ΔG.

The collective variables for adenylate kinase and ribose binding protein were measured with MDAnalysis.^42,43^ For adenylate kinase, the collective variables are the AMP-binding domain and ATP-binding domain angles relative to the core domain. The core domain is composed of residues 1-29, 60-121, and 160-214, the AMP binding domain (Hinge) is residues 30-59, and the ATP binding domain (Lid) is residues 122-159. The collective variables for ribose binding protein define the tilt and twist of the two domains relative to each other.^44,45^ The N-terminal domain is composed of residues 1-100 and 236-259 and the C-terminal domain is composed of residues 108-231 and 269-271. The tilt angle is defined as the angle between the center of mass of the N- and C-terminal domains and the center of mass of the hinge point comprising residues 101-107, 232-235, and 160-268. The twist angle is the dihedral angle of the center of mass of the N-and C-terminal domains and two regions near the ribose binding site on the top of the N-terminal domain, 124-125, 262-262, and 283-284 and the bottom of the C-terminal domain, 133-134, 255-256, 294-295.

## RESULTS AND DISCUSSION

### Energetics of conformational sampling by AlphaFold2

Multiple methods have been advanced to generate conformational ensembles from AlphaFold2, including MSA subsampling methods and the *in silico* mutagenesis method SPEACH_AF.^5,9,10^. An outstanding question is the relationship of the models generated to the energetic landscape of the proteins. We surmised that by evaluating the energetic ranking of these models using the Rosetta Energy Score (RES) function, we could obtain a proxy for the protein conformational landscape. We were stimulated by a previous study, examining the role of mutations in protein stability, that reported the correlation of the change in the RES of a mutant AlphaFold2 model relative to wild-type model (ΔΔG) with the experimental changes in the melting temperature of the protein.^46^

Applying the *in silico* mutagenesis approach of SPEACH_AF and MSA clustering, we characterized the conformational diversity of protein targets and quantified their relative energies. Specifically, we compared the structural diversity and energetics of models generated by the MSA subsampling technique AF_cluster^10^ to the same set of proteins previously shown to be amenable to conformational sampling using SPEACH_AF.^5^ Following generation of models by both methods, a two-step filtering method to parse out misfolded and misthreaded structures was carried out. The first step removed models with a pLDDT less than 70, where a pLDDT of 70 or greater corresponds to a generally correct backbone prediction.^3^ The second step entails principal component analysis (PCA) to identify distorted structures.

Initial tests with AF_cluster, based on pLDDT, yielded very few well-folded models when the minimum number of sequences per cluster was set to 3. Increasing the minimum number of sequences, to either 7 or 11, resulted in a larger percentage of models that passed the two-step filtering, but with a reduction in the total number of models generated (Table 1 and Table S4). For Ribose Binding Protein (RBP) and 3 out of 4 GPCRs (CGRPR, FZD7, and PTH1R), AF_cluster yielded few or no usable models. Table 1 also reports the percentage of models generated by SPEACH_AF that passed the two-step parsing. These remaining models underwent relaxation with Rosetta and were compared to the unrelaxed models using the principal components of the unrelaxed models.

**Table 1:** Initial Screen of AlphaFold2-generated Models.

| Table 1: Initial Screen of AlphaFold2-generated Models<sup>a</sup> |  |  |  |  |
| --- | --- | --- | --- | --- |
| protein | AFC_3<sup>b</sup> | AFC_7<sup>b</sup> | AFC_11<sup>b</sup> | SPEACH_AF |
| AK | 289/2395 (12.1) | 45/305 (14.8) | 11/120 (9.2) | 300/345 (87.0) |
| RBP | 5/1715 (0.3) | 5/330 (1.5) | 5/100 (5) | 420/422 (99.5) |
| MCT1 | 129/1320 (9.8) | 198/360 (55.0) | 169/235 (71.9) | 690/692 (99.7) |
| STP10 | 950/1695 (56.0) | 518/545 (95.0) | 301/305 (98.7) | 735/735 (100) |
| MurJ | 644/1380 (46.7) | 361/445 (81.1) | 199/245 (81.2) | 507/677 (74.9) |
| PfMATE | 588/665 (88.4) | 120/120 (100) | 125/125 (100) | 690/692 (99.7) |
| Lat1 | 486/1780 (27.3) | 429/670 (64.0) | 269/325 (82.8) | 705/705 (100) |
| SERT | 176/1500 (11.7) | 72/570 (12.6) | 154/420 (36.7) | 825/825 (100) |
| ASCT2 | 60/1245 (4.8) | 52/310 (16.8) | 66/130 (50.8) | 690/693 (99.6) |
| ZnT8 | 604/1905 (31.7) | 376/515 (91.7) | 220/240 (91.7) | 450/465 (96.8) |
| CCR5 | 430/1095 (39.3) | 303/435 (69.7) | 221/270 (81.8) | 447/452 (98.9) |
| CGRPR | 0/520 (0) | 0/205 (0) | 3/140 (2.1) | 217/390 (55.6) |
| FZD7 | 10/560 (1.8) | 7/170 (4.1) | 5/105 (4.8) | 508/510 (99.6) |
| PTH1R | 0/105 (0) | 0/50 (0) | 0/35 (0) | 389/452 (86.1) |
| a: Shown are the number of models that passed the initial two-step screen relative to the total number of models generated with the percentage given in parentheses. For a breakdown of the models below 70 pLDDT and distorted (misfolded or misthreaded) see Table S4.<br>b: AFC_3, AFC_7, and AFC_11 utilizing AF_cluster with the minimum number of sequences set to 3, 7, and 11 respectively. |  |  |  |  |

*E. coli* adenylate kinase (AK) is a canonical example of conformational flexibility. AK crystal structures in various catalytic states highlight two flexible domains that independently bind ATP and AMP.^47^ AK models generated by SPEACH_AF and AF_cluster demonstrated little overlap (Fig. 1A). Rosetta relaxation leads to changes in the overlap along PC1 between the two sets of models (Fig. 1B). Evaluation of the models with the relative Rosetta Energy, ΔG, reveals several regions populated with low energy models (Fig. 1C). A plot of the energy vs the first principal component indicates that the lowest energy model is near the closed state, 1AKE, with the next 4 lowest models lying between the closed and fully open states (Fig. 1D-E). Analysis of the ensemble by TM-score and RMSD supports that the AlphaFold2 models sample conformational space spanning the crystal structures (Fig. S1A).

**Figure 1.**
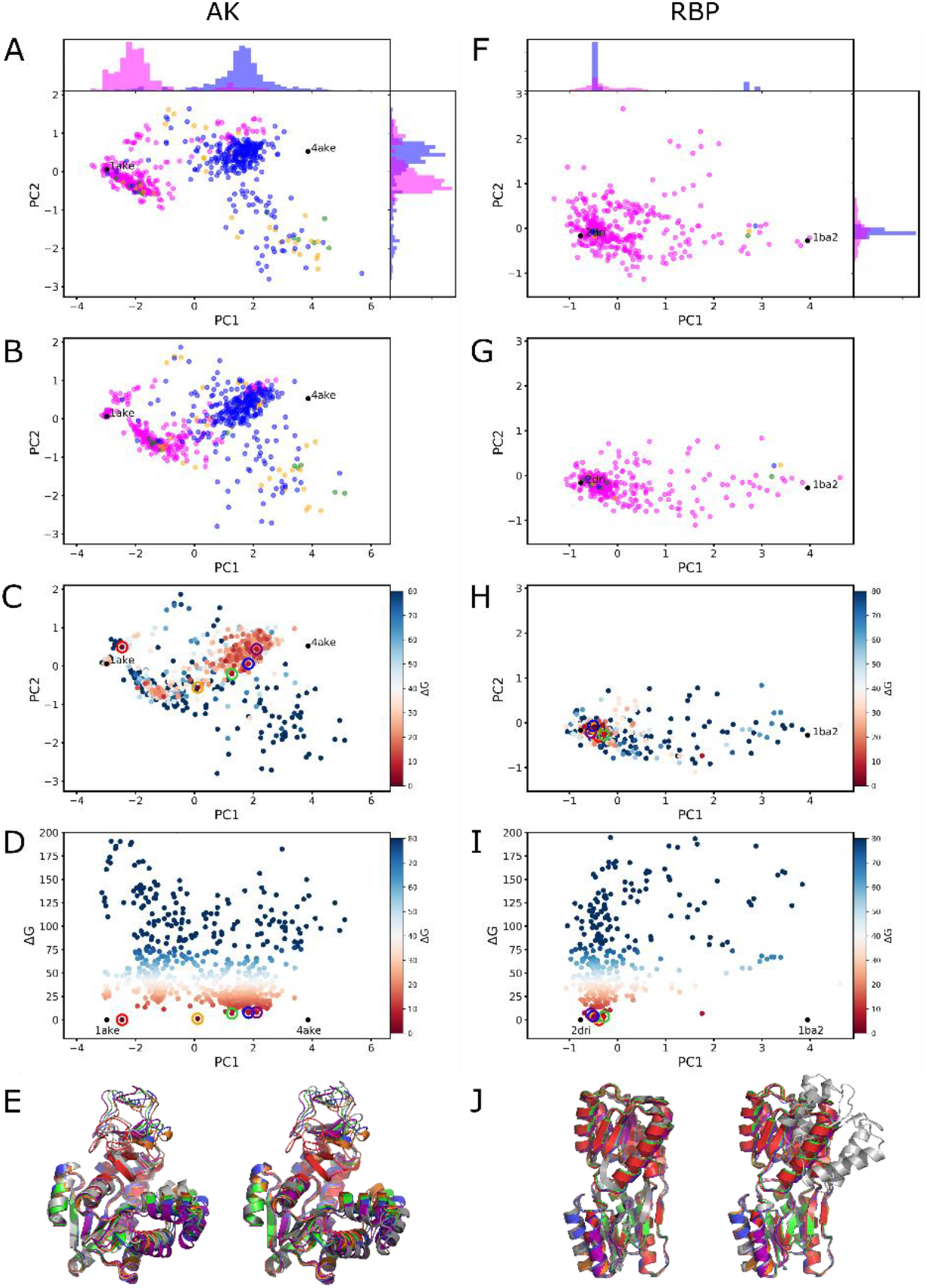
Principal Component Analysis of Adenylate Kinase (AK) and Ribose Binding Protein (RBP). A&F) The first two principal components for the two-step parsed models. The SPEACH_AF models are in magenta, and the AF_cluster models are blue (3), gold (7), and green (11). B&G) The first two principal components for the models after Rosetta relaxation. Colors as in A&F. C&H) Same plots as in B&G except with the coloring based on relative Rosetta Energy Score, ΔG. The five lowest energy models in order, red, orange, green, blue, and purple, are circled. D&I) Plot of ΔG vs PC1. The five lowest energy models, same as in C&H, in order, red, orange, green, blue, and purple, are circled. The crystal structures were not energy minimized and are placed at zero ΔG and their PC1 position for comparison. E) Five lowest energy models shown with the structure from 1ake (left, grey) and 4ake (right, grey). The RMSD for the five models in order from lowest to highest RES to 1ake are 1.96, 3.91, 4.81, 5.20, and 5.53 and to 4ake are 6.91, 4.38, 3.17, 2.76, and 2.60. These values follow the relationship in PC1 space for the models relative to the crystal structures. J) Five lowest energy models shown with the structure from 2dri (left, grey) and 1ba2 (right, grey). The RMSD for the five models in order from lowest to highest RES to 1ba2 are 4.76, 4.86, 4.66, 4.92, and 4.91 and to 2dri are 1.25, 1.20, 1.20, 1.21, and 1.08. These values correlate with the models clustered near the 2dri crystal structure.

If the movements of the ATP lid domain (Lid) and the AMP hinge domain (Hinge) relative to the core domain are independent, there would be four main conformations of AK: Lid closed/Hinge closed, Lid open/Hinge closed, Lid closed/Hinge open, and Lid closed/Hinge closed.^48–50^ We calculated the angle of the Lid and Hinge domain relative to the core domain (Fig. S2A). This plot of collective variables is similar to the plot of the two principal components (Fig. 1B) establishing that the PCA is reflective of these domain movements. An overlay of the AlphaFold2 models on a previously calculated free energy landscape of AK^50^ (Fig. S2B and Fig. S2C) shows that a subset of the models clusters in the low energy well with a Lid angle around 130°, with most of the models lying between this well and the fully open conformation. Remarkably, the conformational pathway of these models follows the trajectories from simulations using dynamic importance sampling method (DIMS) of these conformational changes.^50^ The similarity of the models and molecular dynamic simulations highlights the robustness of this approach. Identifying the energetics of the conformations, both low points and saddle points along conformational space, would aid in picking seeds for downstream molecular dynamic simulations and should allow for a decrease in computational time as was shown for identifying cryptic drug binding sites in AlphaFold2 models.^51^

Unlike AK, modeling of RBP with AF_cluster generated few models that passed the two-step filtering (Table 1). Rosetta relaxation changes the distribution of the models along the principal components (Fig. 1F-G). This decrease in explored conformational space by the Rosetta minimized models appears to arise structurally from a decrease in the twist of the two domains of RBP relative to each other. Similar to AK, the TM-scores and RMSD of the AlphaFold2 models supports that the models sample conformational space spanning the crystal structures (Fig. S1B). Most of the low energy models cluster near the closed conformation (Fig. 1H), although there are a few lower energy models midway between the closed and open conformations (Fig. 1I-J). Parametrization of the AlphaFold2 models with known collective variables, the tilt and twist angle of the two domains relative to each other^44,45^, indicate that the two collective variables are interdependent reflecting coupled conformational changes (Fig. S3). The linear distribution of AlphaFold2 models between the closed (holo) and open (apo) structures is consistent with the trajectories from molecular dynamic simulations.^44,45^

To further establish the robustness of this approach, we expanded our analysis to include benchmark transporters and receptors used in previous examinations of protein conformational diversity generated by AlphaFold2.^5,9^ The conformational space explored by the AF_cluster and SPEACH_AF models differ for the two MFS transporters, MCT1 and STP10. For MCT1, the models span the range between the two crystal structures (Fig. S4A-B and Fig. S5A). The SPEACH_AF models tend to segregate near the two crystal structures whereas the AF_cluster models fill the space between the two crystal structures. There are models of similar low energy across the entire conformational space defined by PC1 (Fig. S4C-D). It is unclear if the low energy models in between the two structures are distinct energy wells in conformational space or distinct points because of under-sampling the conformational space. In contrast, there is a clearer segregation of the models generated by AF_cluster and SPEACH_AF for STP10 (Fig. S4A-B and Fig. S5B). The SPEACH_AF models span PC1 between the two crystal structures whereas the AF_cluster models span a more diffuse conformational space. The low energy models reside between the two crystal structures (Fig. S4C-D).

For the multidrug/oligosaccharidyl-lipid/polysaccharide (MOP) transporter MurJ, there is overlap between the conformations generated by AF_cluster and SPEACH_AF (Fig. S4A-B and Fig. S6A). The low energy models are all in a single cluster slightly biased towards the outward-open conformation, 6NC9 (Fig. S4C-D). Conversely, for PfMATE, which is a member of the same MOP superfamily, there is almost no overlap between the models generated by AF_cluster and SPEACH_AF (Fig. S4A-B and Fig. S6B). The low energy models span most of the conformational space from the outward-open crystal structure, 3VVN, to the inward-open structure, 6FHZ. However, the lowest energy models adopt a more occluded conformation (Fig. S4C-D).

As in most cases, the AF_cluster and SPEACH_AF models have almost no overlap for the two LeuT-fold transporters, LAT1 and SERT (Fig. S7A-B and Fig. S8A-B). For LAT1, the SPEACH_AF models are in a tighter cluster spanning the range between the two crystal structures, while the AF_cluster models span a more diffuse conformational space between the two crystal structures. Examining the energetics of the models for LAT1, we find one main cluster near the outward-open structure, 7DSQ, with a few slightly higher energy models closer to the inward-open structure, 6IRS (Fig. S7C-D). For SERT, the configuration of the AF_cluster and SPEACH_AF models are different compared to LAT1, as they span the conformational space between the two crystal structures, but in a continuous nature with little overlap (Fig. S7A-B and Fig. S8A-B). Similar to LAT1, the low energy models all cluster near the outward-open structure, 5I6X (Fig. S7C-D).

ASCT2 is a sodium-dependent exchanger of neutral amino acids that is thought to work via an elevator-type mechanism. There are very few conformations generated by AF_cluster that pass the two-step filtering method, but in this case the models appear to overlap with the SPEACH_AF models (Fig. S7A-B and Fig. S9A). Most of the AlphaFold2 models align with one of the two the two crystal structures with few models in an intermediate conformation. The low energy models cluster near the 6RVX structure, where the elevator is in the up position.

ZnT8 is a zinc transporter that is a member of the cation diffusion facilitator family. Here again, the AF_cluster and SPEACH_AF models have little overlap. Interestingly, the SPEACH_AF models form a structural progression across the conformational space, but it is the AF_cluster models that collect near one of the crystal structures, 6XPF_A (Fig. S7A-B and Fig. S9B). The low energy models span the conformational space from one crystal structure to the other.

Of the four G-protein coupled receptor (GPCR) proteins in the benchmark set, few or no viable models are generated with AF_cluster for three of the proteins (Table 1). One of these, CGRPR, also had a lower percentage of viable SPEACH_AF models compared to the rest of the benchmark set. We speculate that the information content of the MSA may contribute to the sparsity of models that survive the two-step filtering. Methods to optimize the information content of the MSA for AlphaFold2 predictions are still an intense are of inquiry.^52^ Similarly, the AF_cluster models of CCR5 have little overlap with the SPEACH_AF models (Fig. S10A and Fig. S11A) and relaxation of these models with Rosetta leads to only minor structural changes (Fig. S10B and Fig. S11A). For the other three GPCRs there are not enough AF_cluster models to compare with the SPEACH_AF models. The pattern of low energy models relative to the experimentally determined structures varies across the four GPCRS (Fig. S10C and Fig. S11-S12). This could be due to the conditions that the structures were obtained, such as in the presence of a G-protein or inhibitor, making these conformations only low energy states in their presence.^53^ The inability to sample the full conformational space with the GPCRs is similar to two recent studies examining the ability of AlphaFold2 to generate ensembles for GPCRs.^7,8^ In both studies, diverse models were only obtained by using templates to bias AlphaFold2. This would suggest that while AlphaFold2 has learned elements of the protein energy landscape as encoded in the MSA,^54,55^ it is unable to predict how the landscape is altered in the presence of ligands or accessory proteins.

For most proteins in the benchmark set, the models generated by SPEACH_AF and AF_cluster have little overlap in the conformational space. By mapping the energy scores of the models on the conformational space, the low energy models generally reside in one subset or the other. To further characterize the models, we plotted histograms of the energetics of the models generated by the two methods (Fig. S13-S14). In some cases, there is a clear correlation of one of the methods with the low energy models and this correlation might track with the fold of the protein. For example, SPEACH_AF yielded more low energy models for the LeuT-fold proteins, while for the cation diffusion facilitator ZnT8 AF_cluster models clearly have lower energy. The differential position in conformational space and energetics seen here clearly supports a previous study showing that the MSA content has a role in the models generated by AlphaFold2.^52^ In this study on a method of a differentiable sequence alignment, removal of more diverse sequences lead in one case to better model predictions, but in a second case worse model predictions.

How then does the information content of the MSA reflect on methods that generate models using a single sequence?^56,57^ Could the subsampled MSAs from AF_cluster or modified MSAs from SPEACH_AF highlight how to boost the information content or aid in the learning of the linear sequence for these methods? As we approach the expanded use of AlphaFold2 in structural biology pipelines, further work on how to maximize the information content of the MSA is warranted.^52^ In fact, in CASP 15 two of the methods achieved increased performance over the baseline AlphaFold2 by optimizing the MSA.^58,59^ The question remains how to best create an MSA to yield the breadth of conformations or increase sampling in a computationally tractable way. One method would be to combine AF_cluster with SPEACH_AF as was done with KaiB, where point mutations placed across the MSA could cause the fold switch in KaiB to occur with a single subsampled MSA.^10^

### AlphaFold2 prediction of point mutations

We have been successful in generating alternate conformations by replacing multiple whole columns within the MSA with alanines. In light of these results, we considered the possibility that the effects of the point mutations in AlphaFold2 models in the previous report discussed above^14^ were not observed because the mutation was not placed across the whole MSA. In addition, we considered the possibility that the mutation effects may not be expressed in substantial structural alterations but in changes in the energetics relative to the WT. To test this contention, we compared the two approaches where the mutation was either included in the input sequence only or across all the MSA sequences. We then used structural measures and RES to investigate if Alphafold2 can predict the consequences of the mutations.

Isocyanide hydratase and XylE are two test proteins that undergo conformational changes upon single or double mutations. Isocyanide hydratase and its point mutant D183A, were targets in CASP 15 (T1110/T1109). The mutation induces a change in the active site and a switch in the dimeric structure.^60^ XylE, an MFS transporter, was selected because a double mutation leads to changes in its conformation.^61,62^ Hydrogen/deuterium exchange experiments have shown that only one of the mutations is needed to induce the conformational change.^63^

Modeling of isocyanide hydratase by AlphaFold2 yielded TM scores consistent with accurate prediction of its wild-type structure (Fig. 2A). However, both methods of modeling point mutations, i.e., in the input sequence only and across the MSA, yielded two populations of models. One set of models has a high TM score relative to the wild-type structure (Fig. 2A). Close inspection of these models shows a perturbation around the mutation site, which is reflected by the slightly reduced TM score to the wild-type target (T1110) and a slight increase in the TM-score relative to the mutant target (T1109). The second set of models appears to agree with the structure of the mutant protein (T1109) as reflected in the high TM scores (Fig. 2A and S15B-C). Interestingly, only one of the 5 modelers of AF2 predicted the complete conformational change, but it was not the same modeler in both cases.

**Figure 2.**
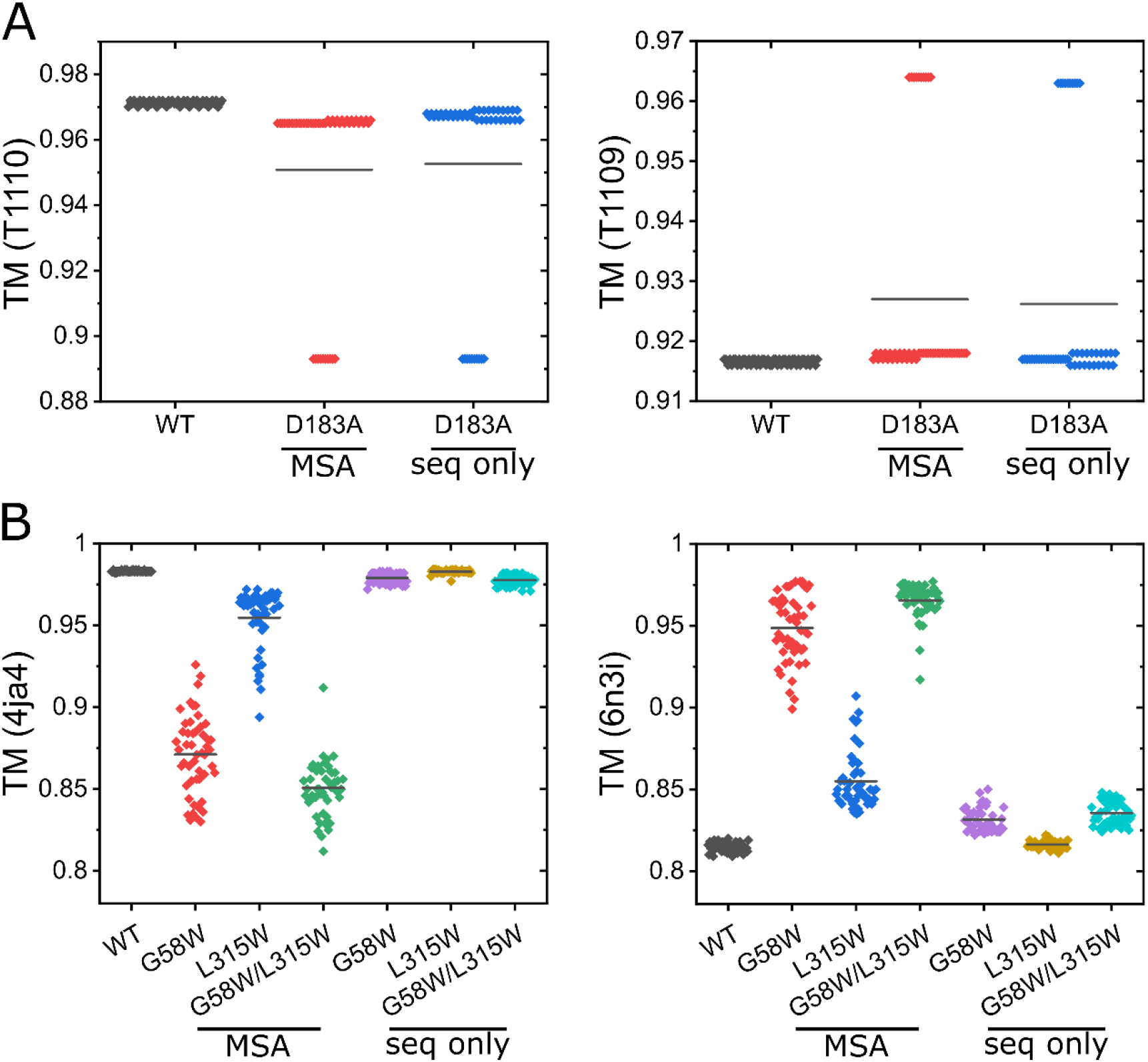
Comparison of WT and Mutant models to crystal structures. A) TM scores for the AlphaFold2 predicted models for wild-type and D183A isocyanide hydratase compared to the target monomers from CASP 15. On the left is the wild-type target (T1110) and on the right is the mutant target (T1109). B) TM scores for the AlphaFold2 predicted models for wild-type and mutant models compared to wild-type (4ja4) and G58W/L315W (6n3i) crystal structures of XylE.

XylE has been crystallized in inward- and outward-facing conformations (Fig. S16A).^61,62^ The outward-facing conformation was stabilized by two point mutations G58A/L315W. Examination of the AlphaFold2 models for XylE indicates that there is little change in TM scores when the mutation is only in the input sequence (Fig. 2B). However, placement of the mutation(s) across the whole MSA improved correlation with the outward-facing state. Both the single, G58W, and double mutant, G58W/L315W, lead to a complete shift in conformation as evidenced by the high TM score when compared to the corresponding experimental conformation of the double mutant (Fig. S16B-C). The G58W mutation also promotes the conformational change in the models in agreement with the hydrogen/deuterium exchange experiments.^63^

Having established that AlphaFold2 can model mutations if the MSA is appropriately modified, we compared the two methodologies on the protein domains used in the Buel and Walters report that concluded that AlphaFold2 is unable to model mutaions.^14^ Their protein set consisted of: the BRCT domain of BRCA1, the ubiquitin-associated domain (UBA) of Rad23 (HR23A), and the MyUb domain of Myosin VI. The mutation A1708E in BRCA1 has been associated with breast cancer^64^ and leads to an increase in proteolytic degradation of BRCA1.^65^ L198A in Rad23 causes disorder of the UBA domain.^66^ The Myosin VI mutation R1117A disrupts the structure of the MyUb domain.^67^

In keeping with the approach of Buel and Walters,^14^ AlphaFold2 models of the mutants were compared to the corresponding domain from the model of the full-length protein from the AlphaFold2 database (Fig. S17). We found that the wild-type BRCT domain of BRAC1 is highly comparable to the domain from the full-length model with an average TM score of 0.986 ± 0.001 (Fig. 3A). Moreover, there is a decrease in the TM-score for the mutant protein regardless of how the mutation is introduced to AlphaFold2. Yet there is a larger change in structure corresponding to a greater average shift in TM-score when the mutation is introduced across the whole MSA (0.974 ± 0.005) compared to having the mutation in the input sequence only (0.976 ± 0.007). The extent of change in TM-scores between each population of mutant models relative to the wild-type is statistically significant in both cases.

**Figure 3.**
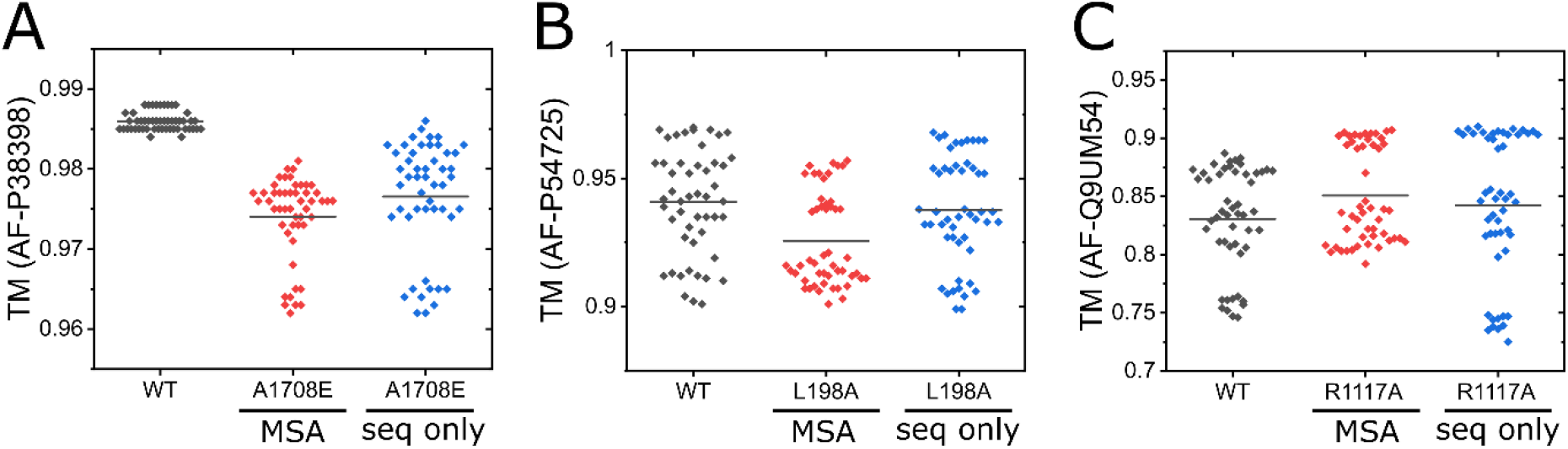
Comparison of WT and Mutant models to reference models. For each of the sets of models a t-test was done to ascertain the confidence that each mutant model set is different from the wild-type model set. A) BRCT domain of BRCA1. The p-values for the t-test and Mann-Whitney U test are 3.55E-22 and 7.05E-18, respectively for in the whole MSA and 1.25E-12 and 8.47E-17, respectively for in the input sequence only. B) UBA domain of HR23A. The p-values for the t-test and Mann-Whitney U test are 1.50E-4 and 4.0E-4, respectively for in the whole MSA and 0.436 and 0.252, respectively for in the input sequence only. C) MyUB domain of Myosin VI. The p-values for the t-test and Mann-Whitney U test are 0.020 and 0.065, respectively for in the whole MSA and 0.284 and 0.159, respectively for in the input sequence only.

The TM-scores for the wild-type UBA, 0.941 ± 0.020, and MyUb, 0.831 ± 0.044, domains modeled in isolation are lower and more diverse than for the wild-type BRCT domain, suggesting that the context of these domains in the full-length protein is important for AlphaFold2 modeling of their structures (Fig. 3B-C). For the UBA domain, there is a smaller change in the TM-score for the models with the mutation in the input sequence, 0.938 ± 0.018, compared to a more significant shift with the mutation across the whole MSA, 0.926 ± 0.021 (Fig. 3B). The TM-scores for the mutant MyUb domain are slightly shifted compared to the scores for wild-type models, though the effects are different depending on the placement of the mutation in the input sequence only, 0.842 ± 0.061, or across the whole MSA, 0.851 ± 0.040 (Fig. 3C). This shift is significant only when the mutation is in the whole MSA.

We next examined whether the RES analysis could provide further insight into the consequences of the mutations for these three proteins. Shown in Figure S18 are the RESs for the BRCT, UBA, and MyUb domains. The change in average RES for the BRCT domain of BRCA1 is shifted 10 units higher for both methods of inputting the mutation. The scale of change in energy suggests that this mutation is destabilizing, potentially driving unfolding. In contrast, the mutants of the UBA domain of HR23A lead to a slight increase in the average value, while the change for the MyUb domain of MyoVI is marginal. In both cases the change in RES is slightly larger for the models predicted from the mutation introduced only in the input sequence. This appears to be contrary to the results for the TM-score, where there was a slightly larger effect when the mutation was in the whole MSA. Although speculative at this point, it is possible that these results reflect that the structures generated by AlphaFold2 with the mutation in the MSA are more structurally perturbed, leading to the attempt by AlphaFold2 to compensate for the mutation, in effect minimizing the energetic consequences of the mutation. Without this minimization, the change in free energy, as calculated by Rosetta, would be expected to increase in the presence of the mutation in the more wild-type structure.

### Broader characterization of AlphaFold2 predictions of point mutations

The results discussed above bring into the forefront the more general question of how mutations affect protein structure. The wealth of experimental studies highlights that while these tend to be protein specific, they can be grouped into two major classes. In one class, the mutations do not lead to major distortion in the native structure but are mostly associated with changes in the free energy of unfolding. If the change is large enough, the protein does not fold, misfolds, or is unstable and unfolds. In the second class, the mutations lead to notable distortions in the native structure. Thus, at the origin of the different conclusions between our results and those reported earlier^14^ is the expectation that AlphaFold2 should not generate a folded structure since experimentally the proteins were reported to be disordered or unfolded. To gain better insight into the predictive ability of this methodology on the effect of point mutations we next explored two proteins with large experimental data sets testing the effect of point mutations.

As noted above, drastic mutations do not necessarily lead to distortion in the structure but can lead to changes in the free energy of unfolding. A notable example that established these principles is the bacteriophage protein T4 lysozyme (T4L) where numerous mutations were characterized by their crystal structures and free energy of unfolding. Thus, to investigate the relationship between the models of mutations and the energetics, we applied the AlphaFold2/Rosetta energy pipeline to a large set of T4L mutants (Table S2). Figure 4A shows a strong correlation, based on the Spearman’s rank correlation, between the calculated RES and the experimental ΔΔG indicating that these models capture changes in residue interactions reflective of the experimental results. In contrast, we found almost no discernable correlation between various measures of structural similarity or deviations between models of the mutant relative to the crystal structure of the WT and ΔΔG (Fig. S19). While there is a weak correlation with overall pLDDT and a moderate correlation with the normalized pLDDT for the mutated residue (see methods), these are not predictive on an individual amino acid basis.

**Figure 4.**
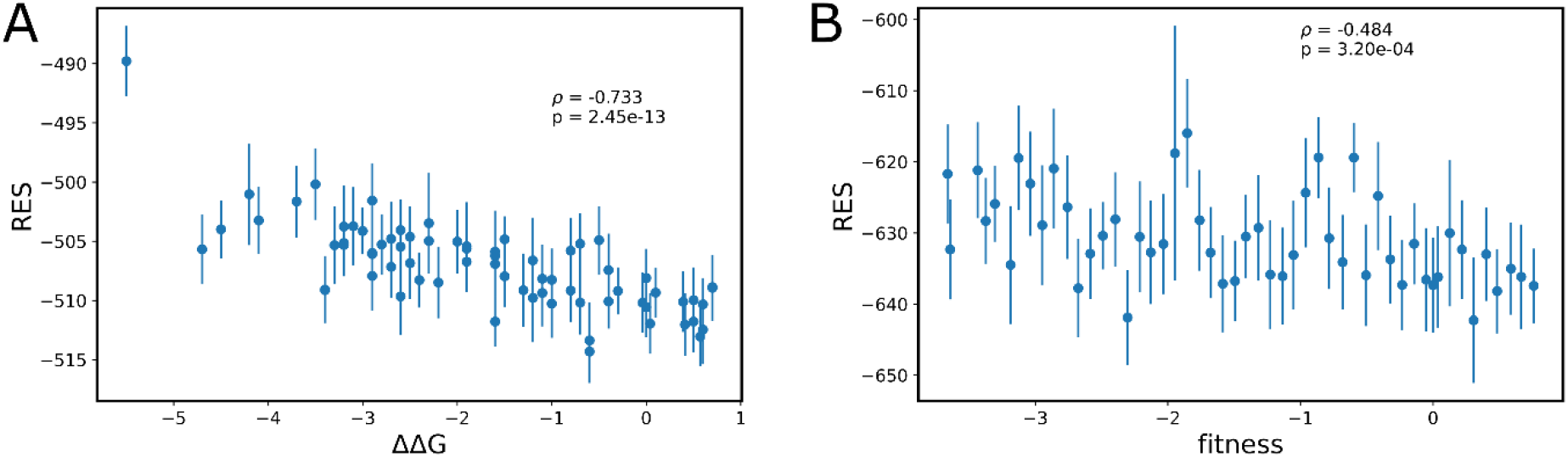
Comparison of Rosetta Energy Scores (RES) to biological readouts for T4 Lysozyme and BRCA1. A) The RES is highly correlative to experimentally determined ΔΔGs with a Spearman’s correlation coefficient of -0.733 and p-value of 2.45E-13. B) The RES is moderately correlative to the fitness measure with a Spearman’s correlation coefficient of -0.48 and p-value of 3.2E-4.

Another common assessment of mutational effects is the fitness score. Indeed, these rapid readouts of large mutational screens provide a facile way to identify hot spots in protein targets. To test the AlphaFold2/Rosetta pipeline, we selected 50 mutations in the BRCT domain, included in the ProteinGym dataset, spanning the range between the highest and lowest fitness values (Table S3).^38,39^ The RES of these models shows moderate correlation to the fitness score (Fig. 4B). Comparing RES for the mutant models to the AlphaMissense score^68^, a metric describing the pathogenicity of single point mutations based on the AlphaFold2 network, indicates a similar correlation as to the fitness score (Fig. S20A). This is expected with the high correlation between the fitness and AlphaMissense scores (Fig. S20B), though there are some outliers in the lower left corner where both the fitness and AlphaMissense scores are low (A1823S, A1823P, A1823T, and P1812A). Understanding the outliers, while beyond the scope of this work, is central to improving AlphaMissense or other predictors of mutations to be applicable for clinical use. There is a confident, weak correlation between the TM-score or RMSD with the fitness value, though the absolute change in values for the TM-score and RMSD are exceedingly small, < 0.05 and 0.1, respectively (Fig. S20C-D). There is a weak correlation with overall pLDDT and little to no correlation with the normalized pLDDT for the mutated residue (Fig. S20E-F). While there is a ranked correlation between the different structural metrics and the fitness score for the mutants, the change is not predictive on an individual mutant basis.

Together, these results would suggest then that AlphaFold2 can predict changes due to mutations, though the structural or energetic consequences of that mutation might not be accurately predicted, especially in the case of mutations that lead to folding defects where repacking by AlphaFold2 may be a consideration. Overall, our analysis supports the conclusion that AlphaFold2 has learned about the folded protein landscape that is encoded in the MSA.^54,55^ Therefore, the parameterization of the energetic and/or structural change is critical when testing the ability of AlphaFold2 to predict the effect of mutations.

## CONCLUSIONS

While AF_cluster was shown to be able to generate both conformations for fold-switch proteins, we find that in some cases it can generate conformational ensembles that are complementary to those generated with SPEACH_AF. The Rosetta scoring of these models effectively maps out energy landscapes that track with previously reported conformations and energetics. We have also shown that AlphaFold2 can generate models that are sensitive to point mutations. The structural changes in the models are more robust when the mutation(s) are in the whole MSA versus just in the input sequence. The combined results support that AlphaFold2 has learned information about protein energy landscapes from information encoded in the MSAs and the structures that it was trained on.

## ACKNOWLEDGEMENTS

We thank Derek Claxton for many helpful discussions and critically reading the manuscript. This work was supported by the National Institute of Health grants GM128087 and GM077659 awarded to HSM.

---

**Fig. S1.**
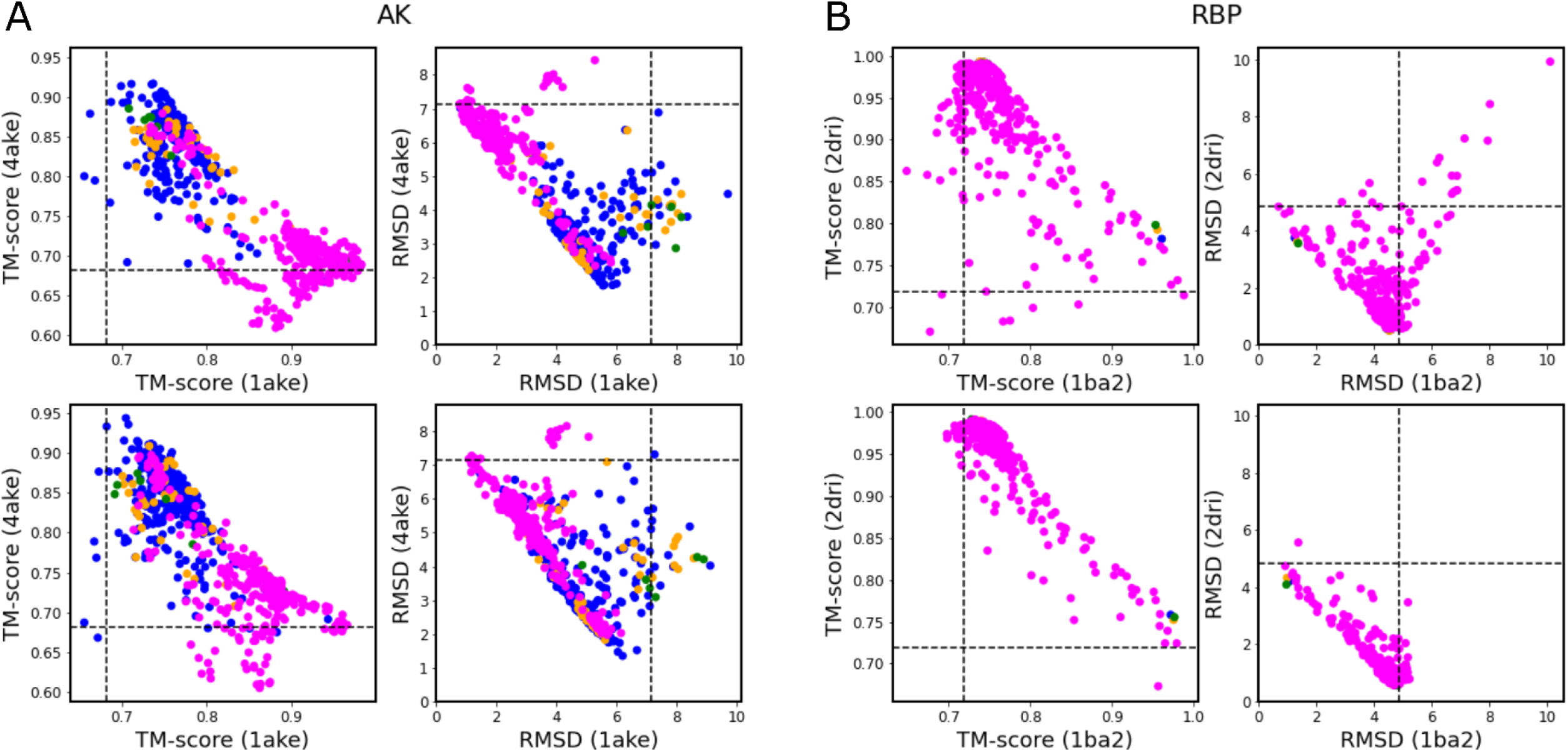
TM-score and RMSD Analysis. Plots of the TM-score against the two experimental structures for Adenylate Kinase (A) and Ribose Binding Protein (B). The plots for the AlphaFold2 models before Rosetta minimization are on top and the models after minimization are on the bottom. The SPEACH_AF models are in magenta, and the AF_cluster models are blue (3), gold (7), and green (11).

**Figure S2.**
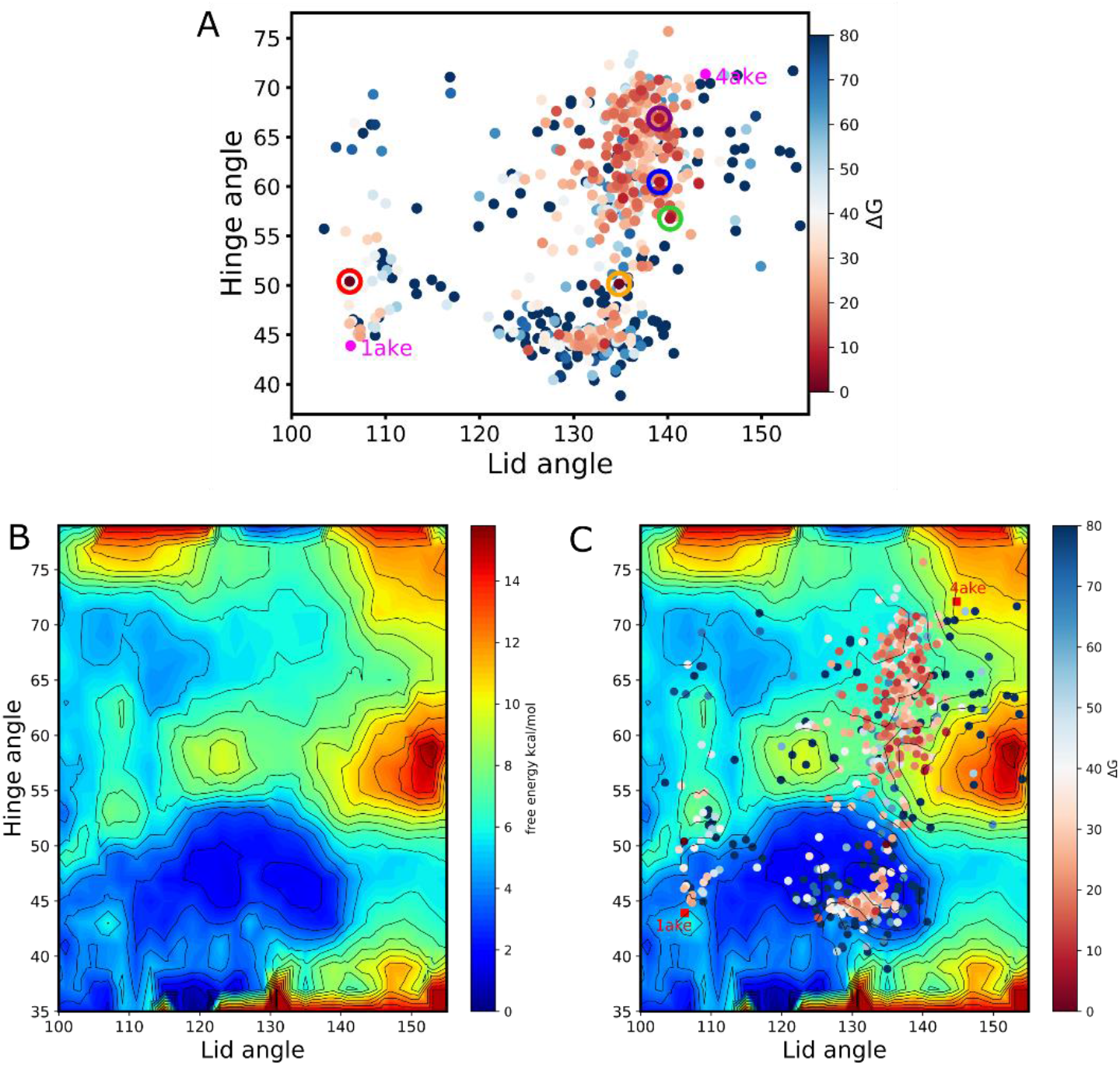
Domain movements in Adenylate Kinase. A) Plot of the angle between the Lid (ATP binding) domain and Hinge (AMP binding) domain to the core domain. The core domain is comprised of residues 1-29, 60-121, and 160-214; the hinge domain is residues 30-59; and the lid domain is residues 122-159. The coloring of this plot corresponds to Fig. 3C. B) Previously published free energy landscape for the Lid and Hinge domain angles with the core domain. C) Overlay of the AlphaFold2 models on the free energy landscape. Pathway of the AlphaFold2 models from the open conformation, 4ake, to the closed conformation, 1ake, follows the trajectories from DIMS simulations of the conformational change. Free energy landscape available from https://becksteinlab.physics.asu.edu/research/52/adk-apo-pmf (O. Beckstein, E. J. Denning, J. R. Perilla, and T. B. Woolf. Zipping and unzipping of adenylate kinase: Atomistic insights into the ensemble of open/closed transitions. J. Mol. Biol., 394(1):160–176, 2009).

**Figure S3.**
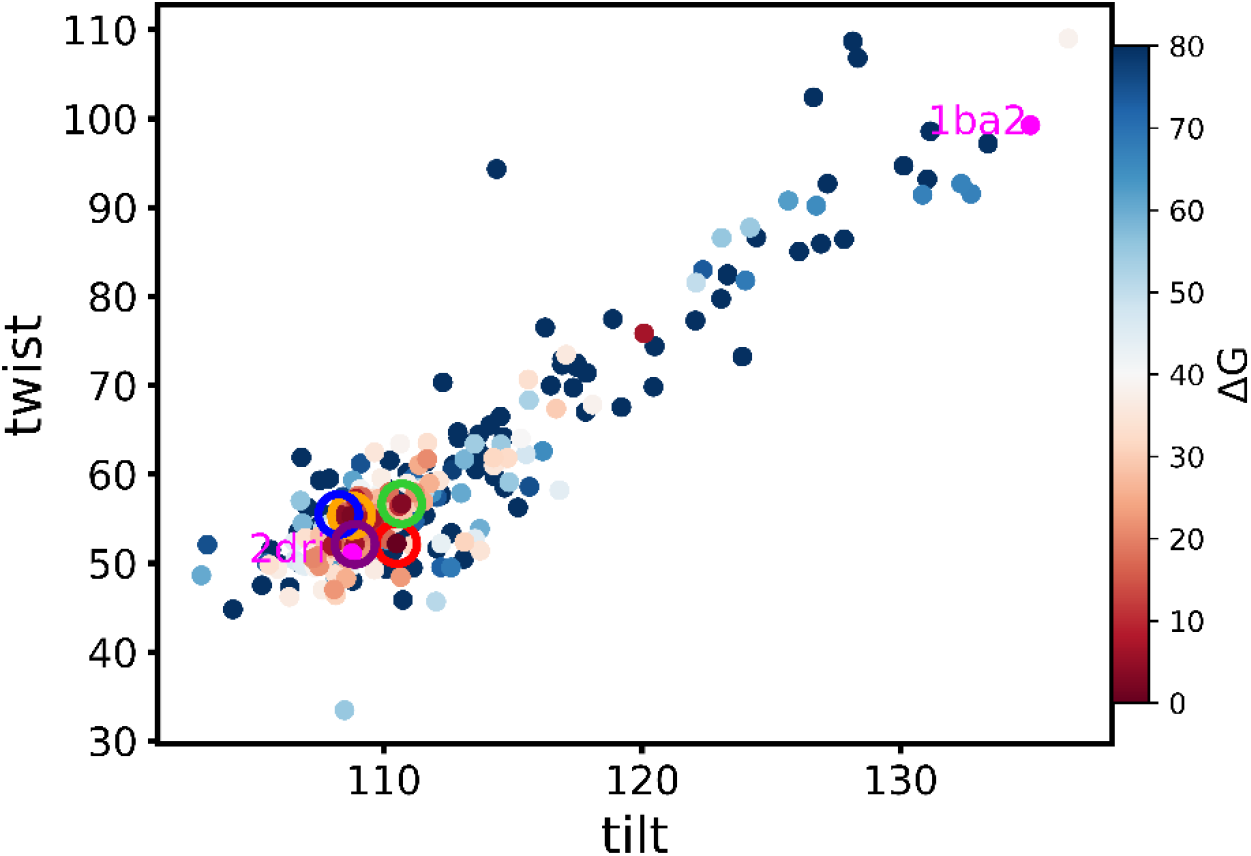
Domain movement in Ribose Binding Protein. Plot of the tilt and twist angle between the N- and C-terminal domains. The N-terminal domain is comprised of residues 1-100 and 236-259 and the C-terminal domain is comprised of residues 108-231 and 269-271. The tilt angle is defined as the angle between the center of mass of the N- and C-terminal domains and the center of mass of the hinge point comprising residues 101-107, 232-235, and 160-268. The twist angle is the dihedral angle of the center of mass of the N- and C-terminal domains and two regions near the ribose binding site on the top of the N-terminal domain, 124-125, 262-262, and 283-284 and the bottom of the C-terminal domain, 133-134, 255-256, 294-295.

**Fig. S4.**
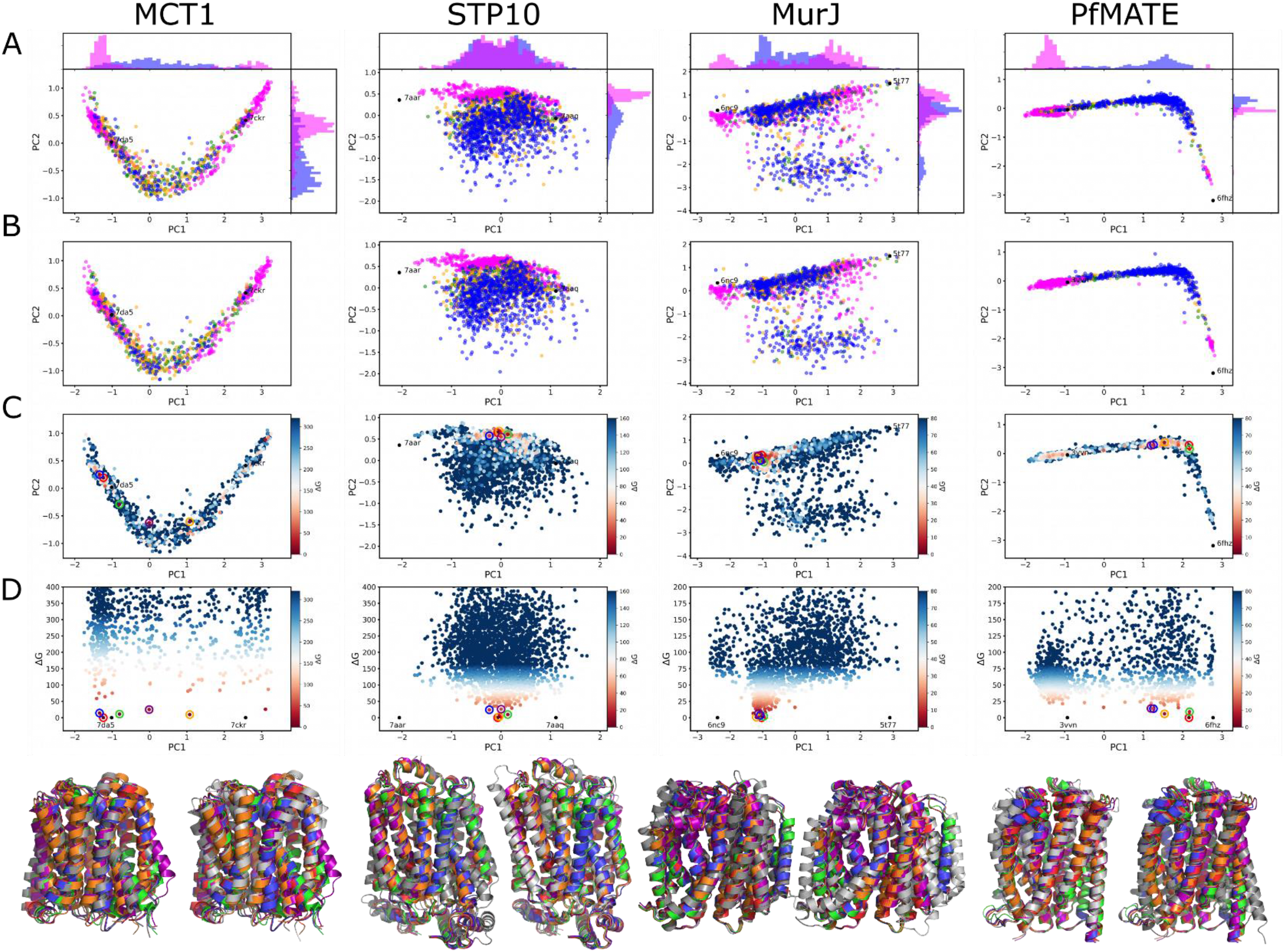
Principal Component Analysis. A) The first two principal components for the two-step parsed models. The SPEACH_AF models are in magenta, and the AF_cluster models are blue (3), gold (7), and green (11). B) The first two principal components for the models after Rosetta relaxation. Colors as in A. C) Same plots as in B except with the coloring based on relative Rosetta Energy Score, ΔG. The five lowest energy models in order, red, orange, green, blue, and purple, are circled. D) Plot of ΔG vs PC1. The five lowest energy models, same as in C, in order, red, orange, green, blue, and purple, are circled. The crystal structures were not energy minimized and are placed at zero ΔG and their PC1 position for comparison. At the bottom are the five lowest energy models shown with the representative structures.

**Fig. S5.**
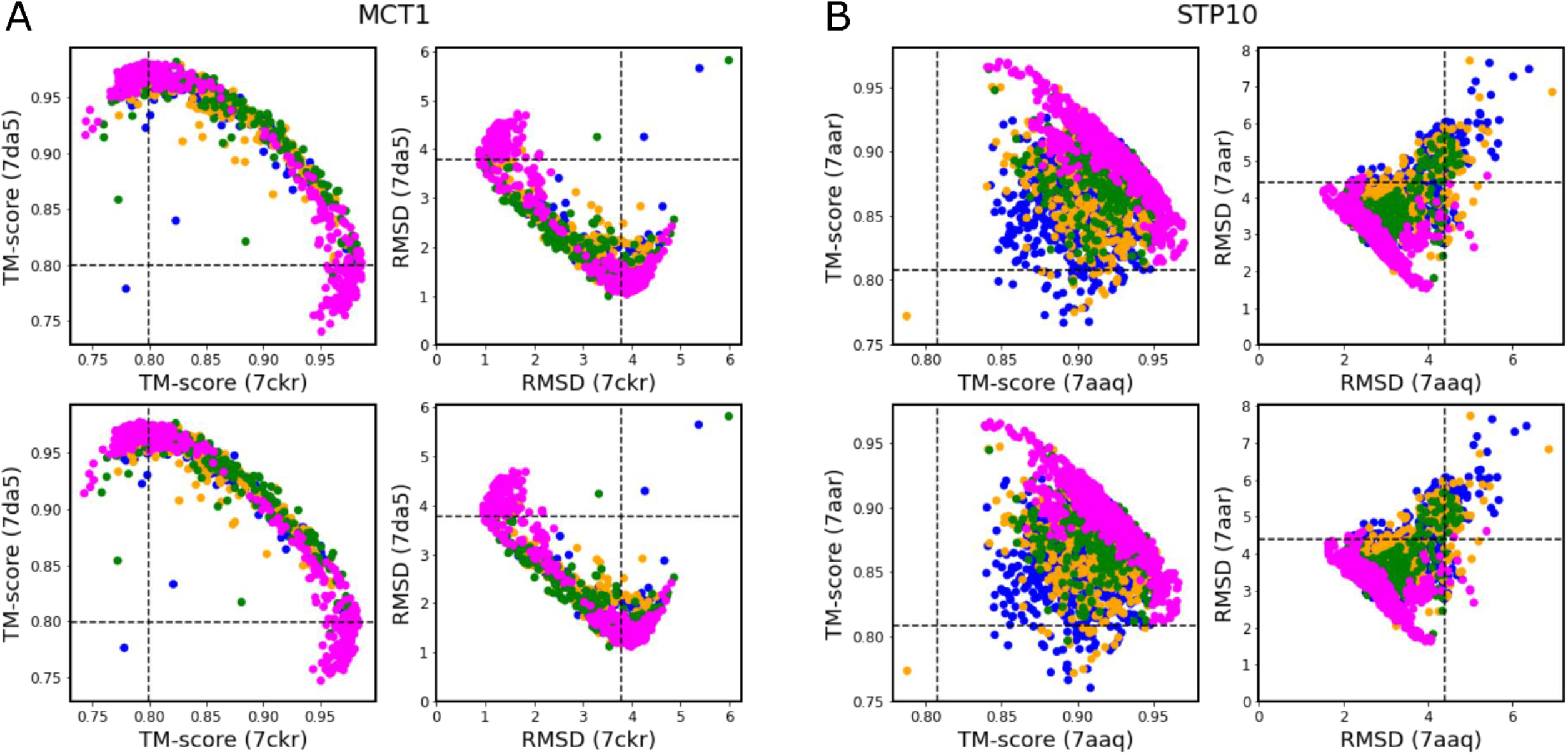
TM-score and RMSD Analysis. Plots of the TM-score against the two experimental structures for MCT1 (A) and STP10 (B). The plots for the AlphaFold2 models before Rosetta minimization are on top and the models after minimization are on the bottom. The SPEACH_AF models are in magenta, and the AF_cluster models are blue (3), gold (7), and green (11).

**Fig. S6.**
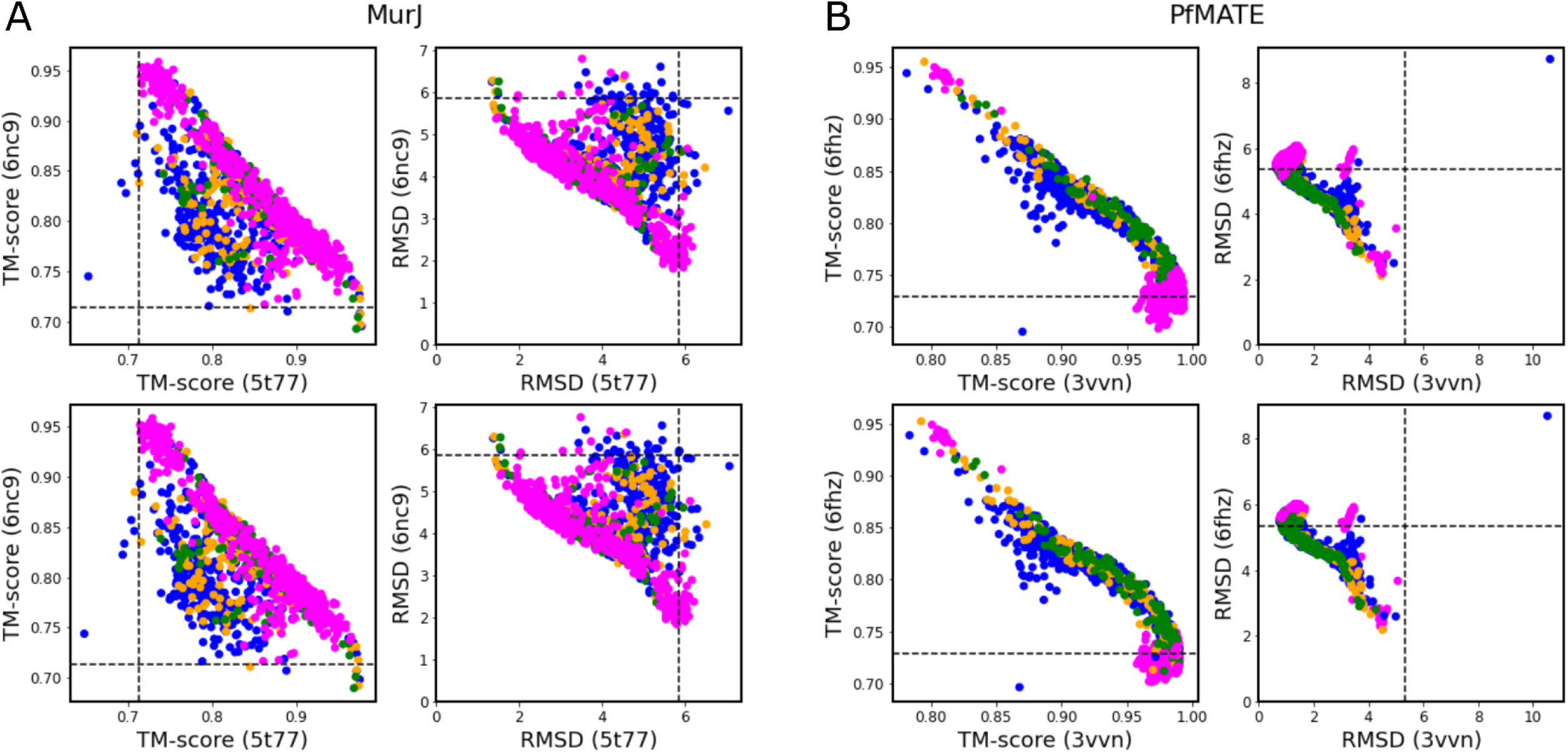
TM-score and RMSD Analysis. Plots of the TM-score against the two experimental structures for MurJ (A) and PfMATE (B). The plots for the AlphaFold2 models before Rosetta minimization are on top and the models after minimization are on the bottom. The SPEACH_AF models are in magenta, and the AF_cluster models are blue (3), gold (7), and green (11).

**Fig. S7.**
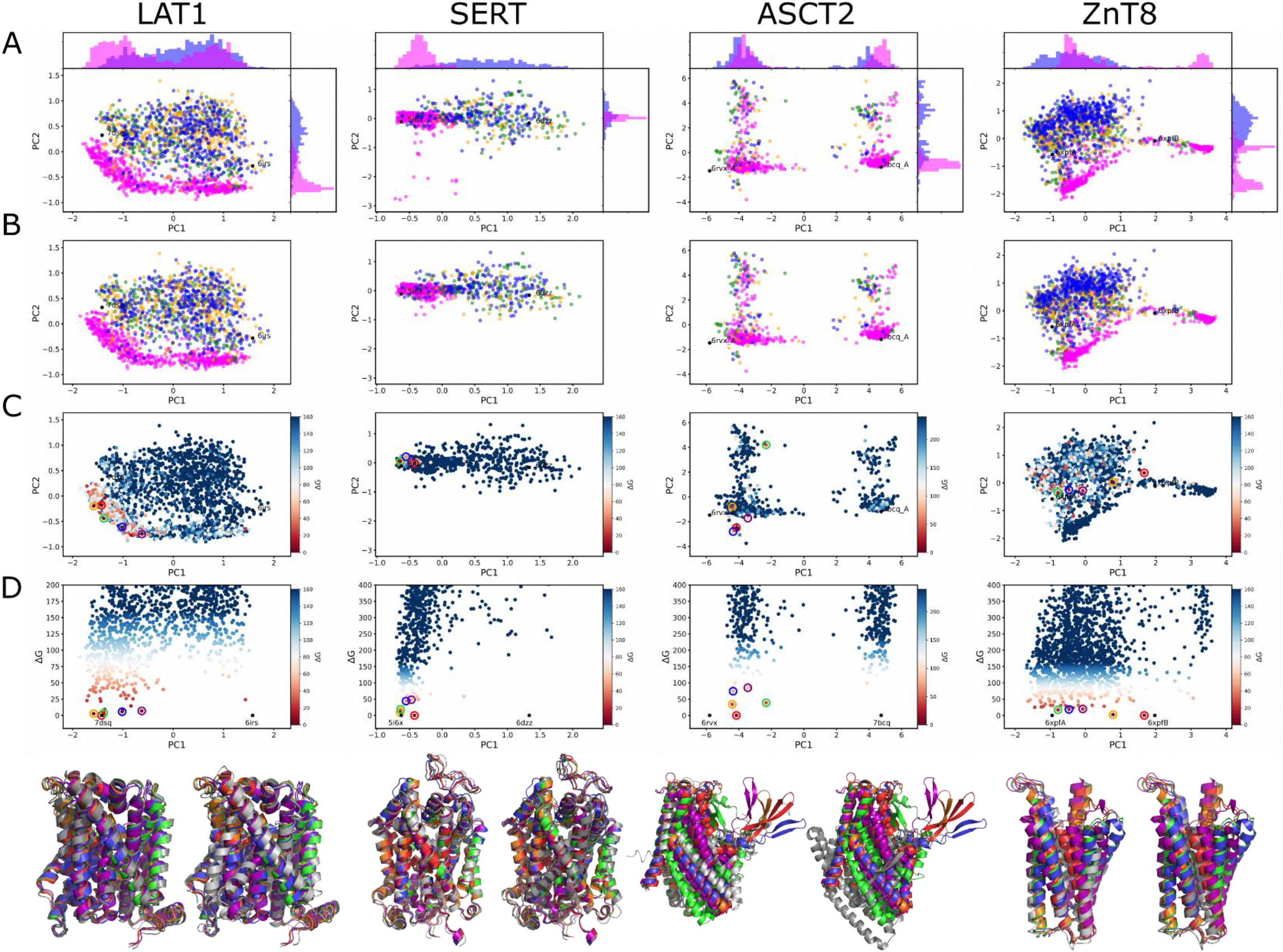
Principal Component Analysis. A) The first two principal components for the two-step parsed models. The SPEACH_AF models are in magenta, and the AF_cluster models are blue (3), gold (7), and green (11). B) The first two principal components for the models after Rosetta relaxation. Colors as in A. C) Same plots as in B except with the coloring based on relative Rosetta Energy Score, ΔG. The five lowest energy models in order, red, orange, green, blue, and purple, are circled. D) Plot of ΔG vs PC1. The five lowest energy models, same as in C, in order, red, orange, green, blue, and purple, are circled. The crystal structures were not energy minimized and are placed at zero ΔG and their PC1 position for comparison. At the bottom are the five lowest energy models shown with the representative structures.

**Fig. S8.**
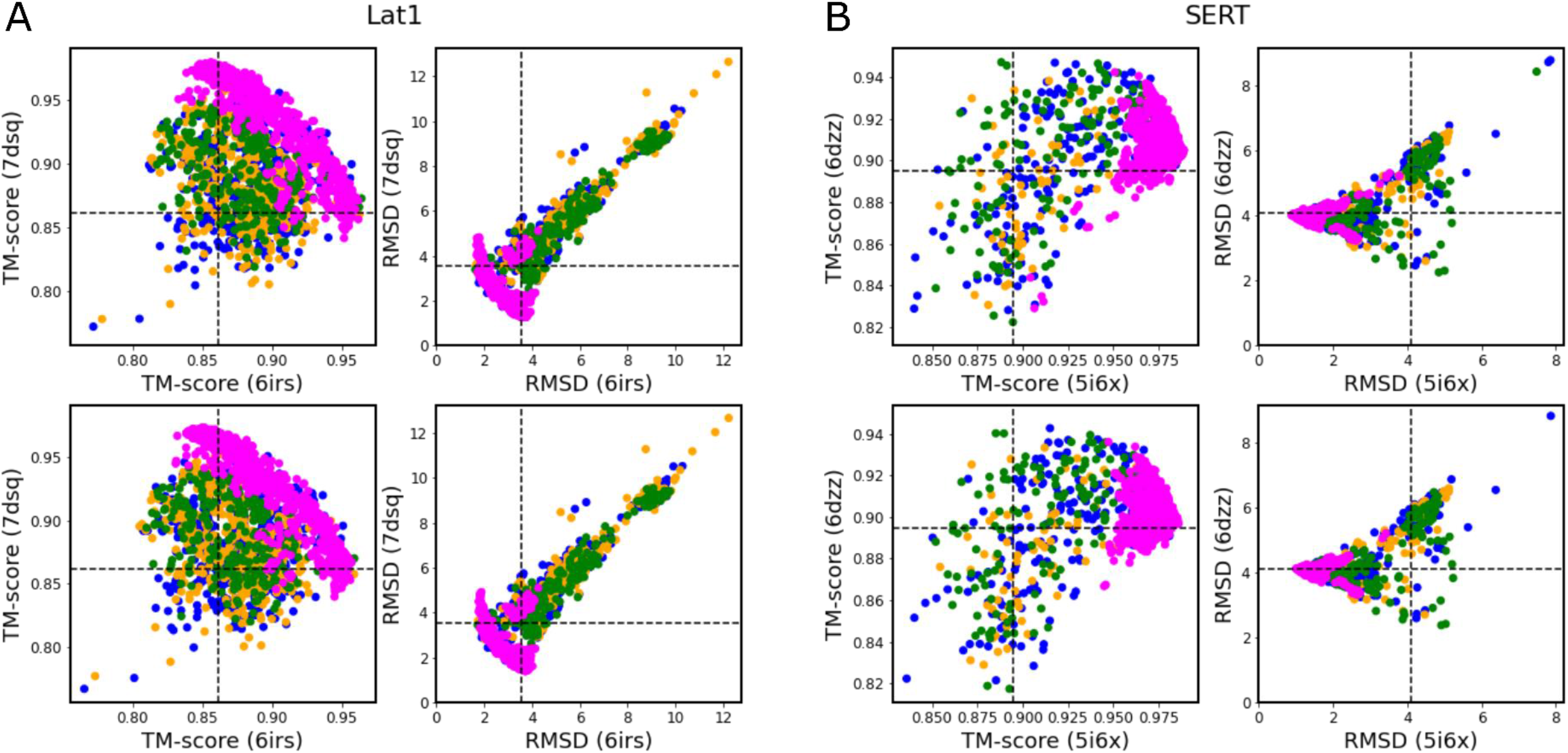
TM-score and RMSD Analysis. Plots of the TM-score against the two experimental structures for LAT1 (A) and SERT (B). The plots for the AlphaFold2 models before Rosetta minimization are on top and the models after minimization are on the bottom. The SPEACH_AF models are in magenta, and the AF_cluster models are blue (3), gold (7), and green (11).

**Fig. S9.**
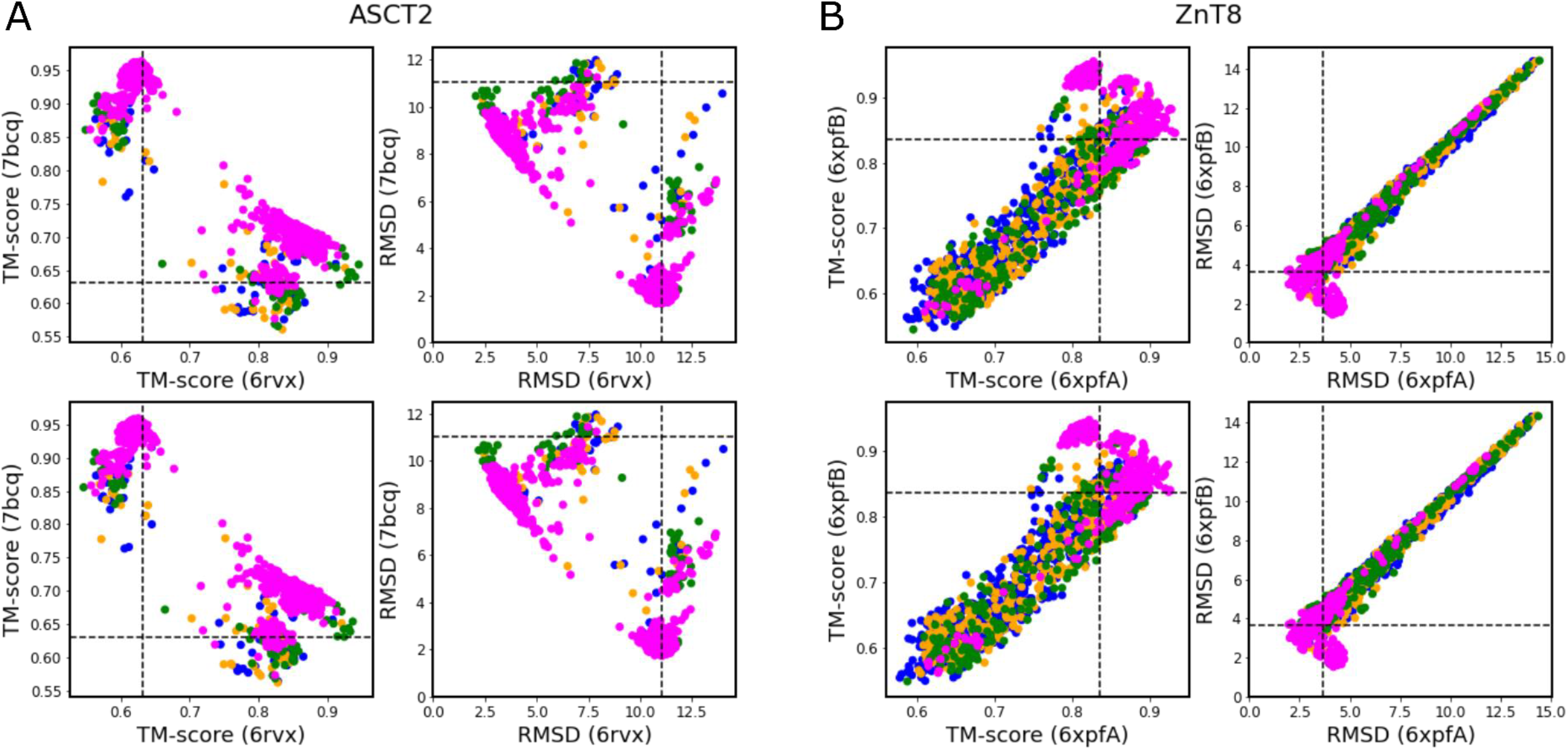
TM-score and RMSD Analysis. Plots of the TM-score against the two experimental structures for ASCT2 (A) and ZnT8 (B). The plots for the AlphaFold2 models before Rosetta minimization are on top and the models after minimization are on the bottom. The SPEACH_AF models are in magenta, and the AF_cluster models are blue (3), gold (7), and green (11).

**Fig. S10.**
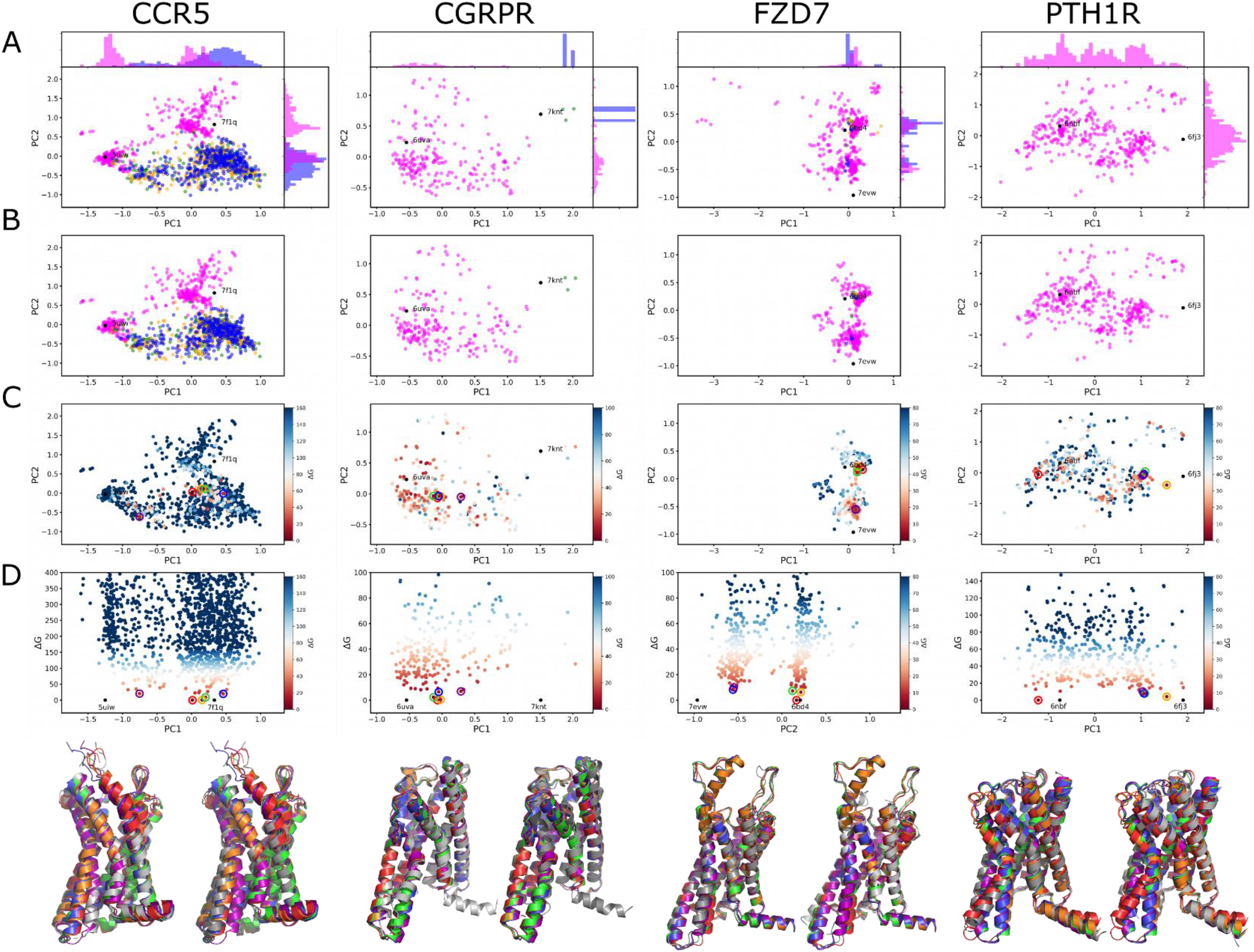
Principal Component Analysis. A) The first two principal components for the two-step parsed models. The SPEACH_AF models are in magenta, and the AF_cluster models are blue (3), gold (7), and green (11). B) The first two principal components for the models after Rosetta relaxation. Colors as in A. C) Same plots as in B except with the coloring based on relative Rosetta Energy Score, ΔG. The five lowest energy models in order, red, orange, green, blue, and purple, are circled. D) Plot of ΔG vs PC1. The five lowest energy models, same as in C, in order, red, orange, green, blue, and purple, are circled. The crystal structures were not energy minimized and are placed at zero ΔG and their PC1 position for comparison. At the bottom are the five lowest energy models shown with the representative structures.

**Fig. S11.**
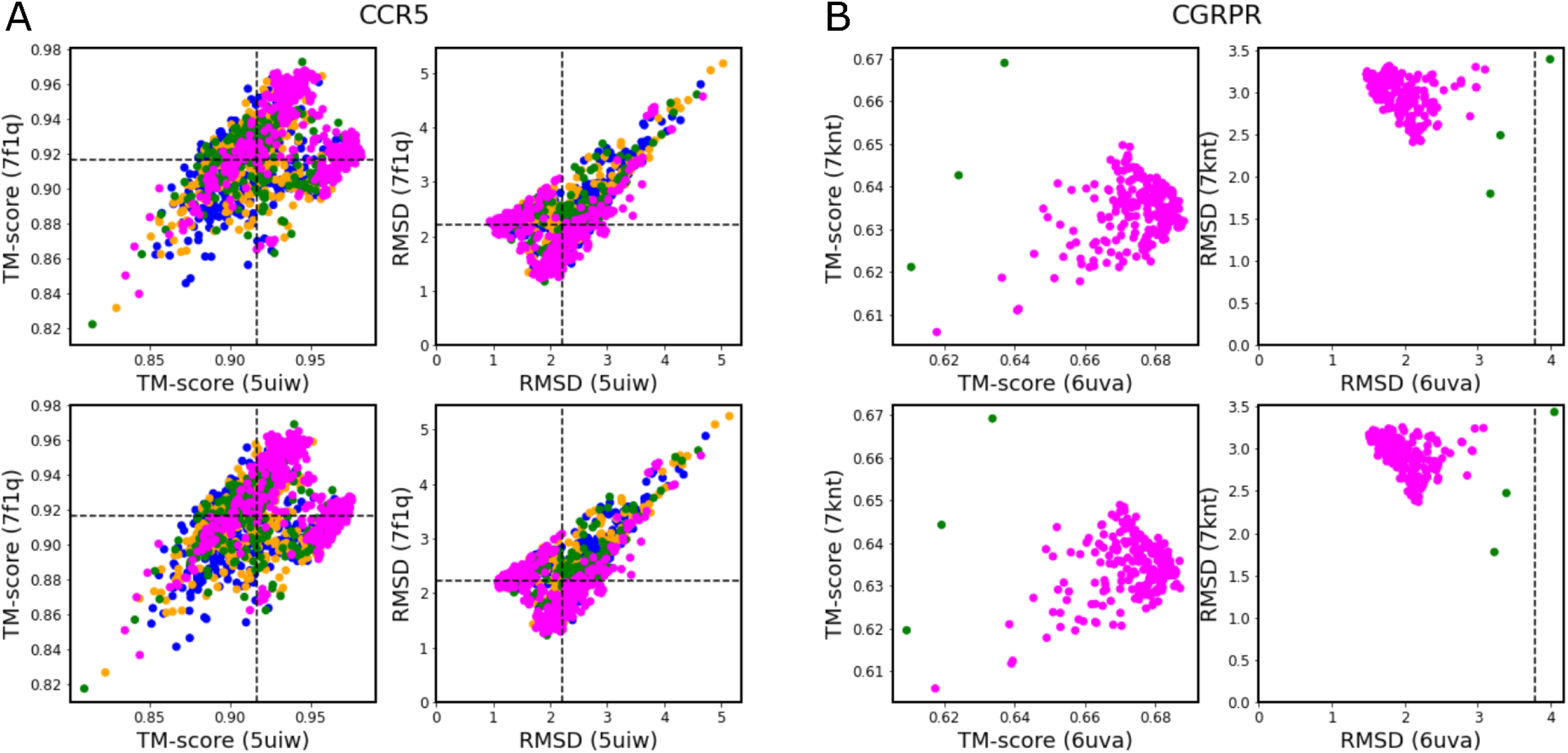
TM-score and RMSD Analysis. Plots of the TM-score against the two experimental structures for CCR5 (A) and CGRPR (B). The plots for the AlphaFold2 models before Rosetta minimization are on top and the models after minimization are on the bottom. The SPEACH_AF models are in magenta, and the AF_cluster models are blue (3), gold (7), and green (11).

**Fig. S12.**
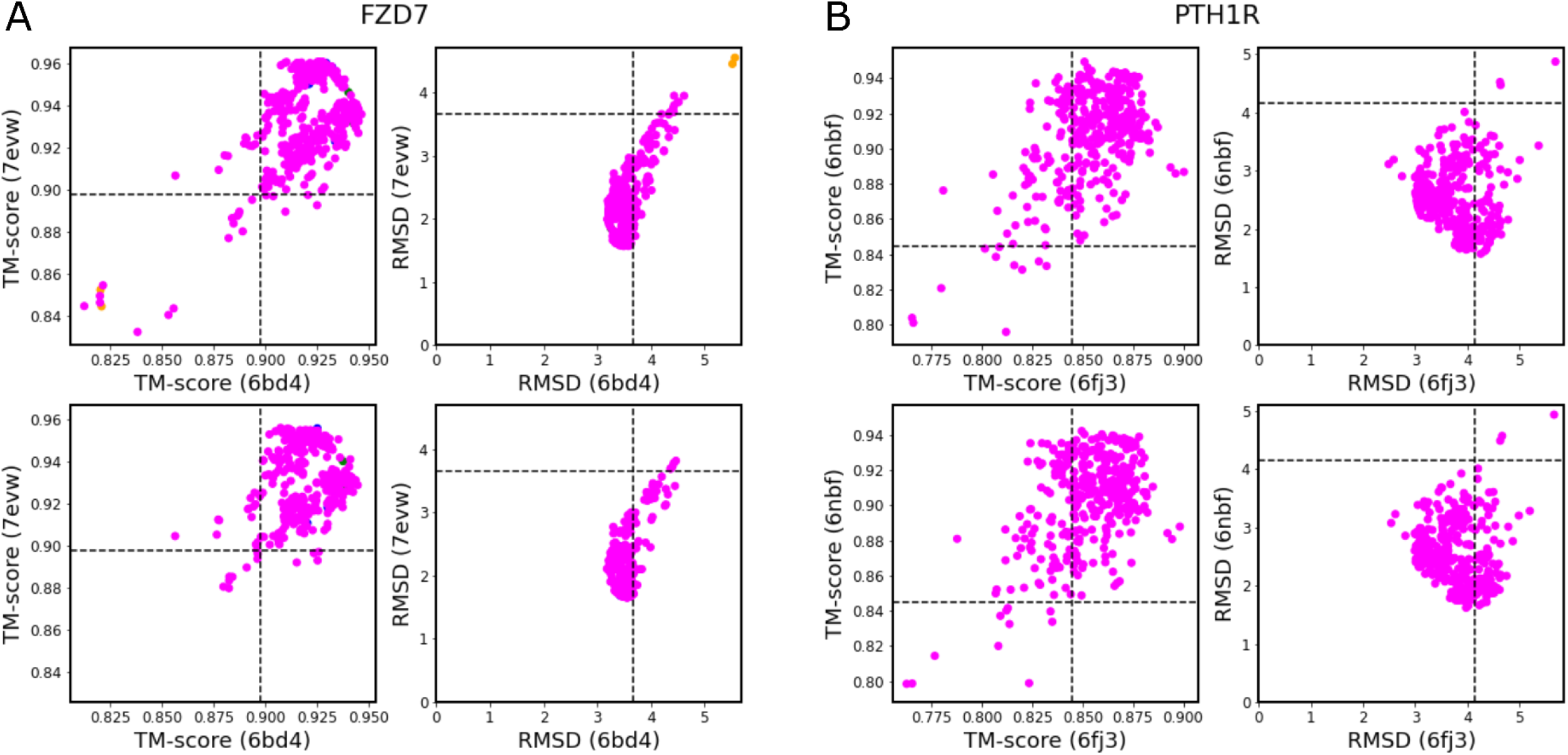
TM-score and RMSD Analysis. Plots of the TM-score against the two experimental structures for FZD7 (A) and PTH1R (B). The plots for the AlphaFold2 models before Rosetta minimization are on top and the models after minimization are on the bottom. The SPEACH_AF models are in magenta, and the AF_cluster models are blue (3), gold (7), and green (11).

**Fig. S13.**
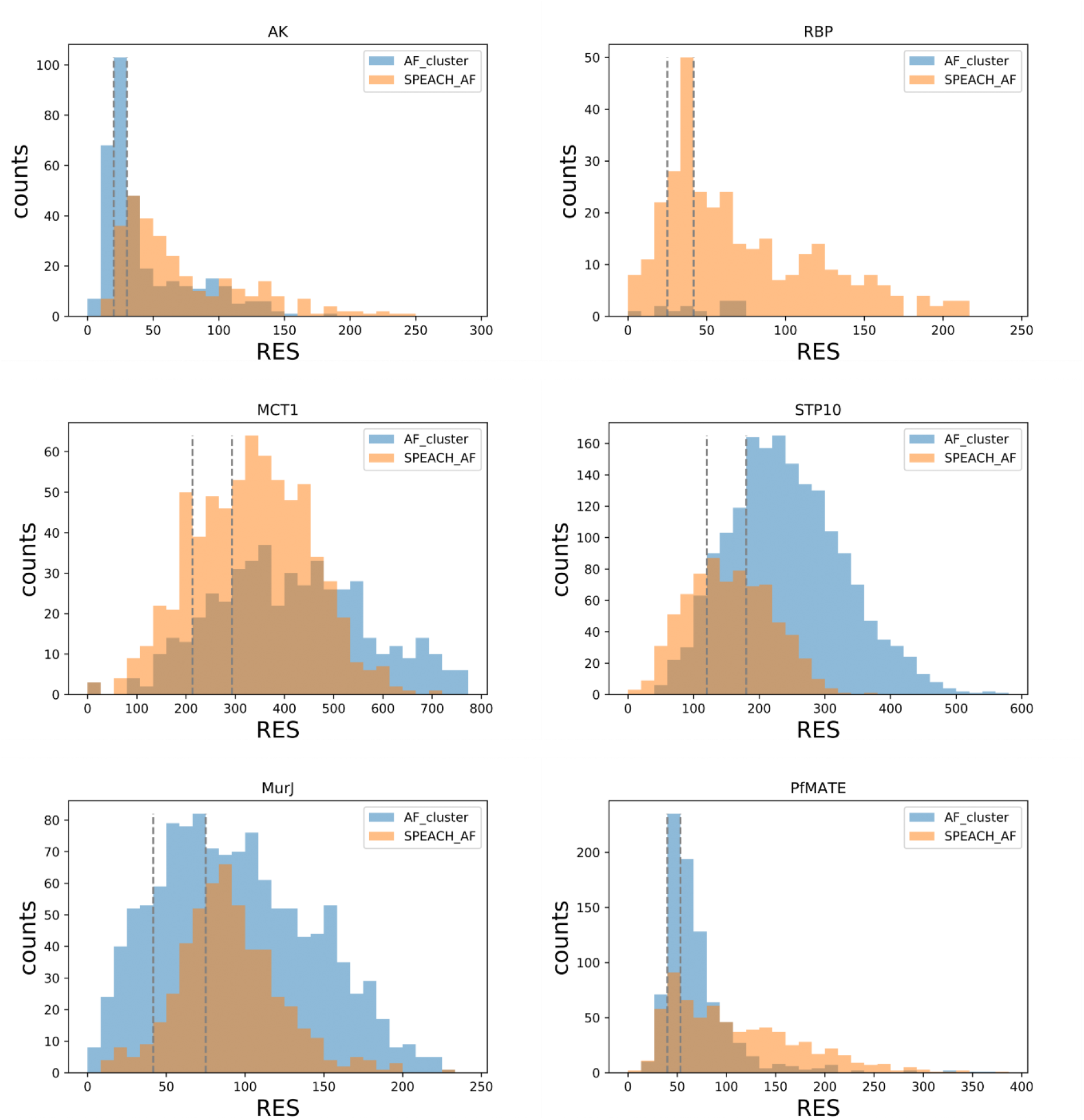
Histograms of RES for AF_cluster and SPEACH_AF. The gray dashed lines demarcate the ∼10% and ∼30% lowest scoring models from both methods combined.

**Fig. S14.**
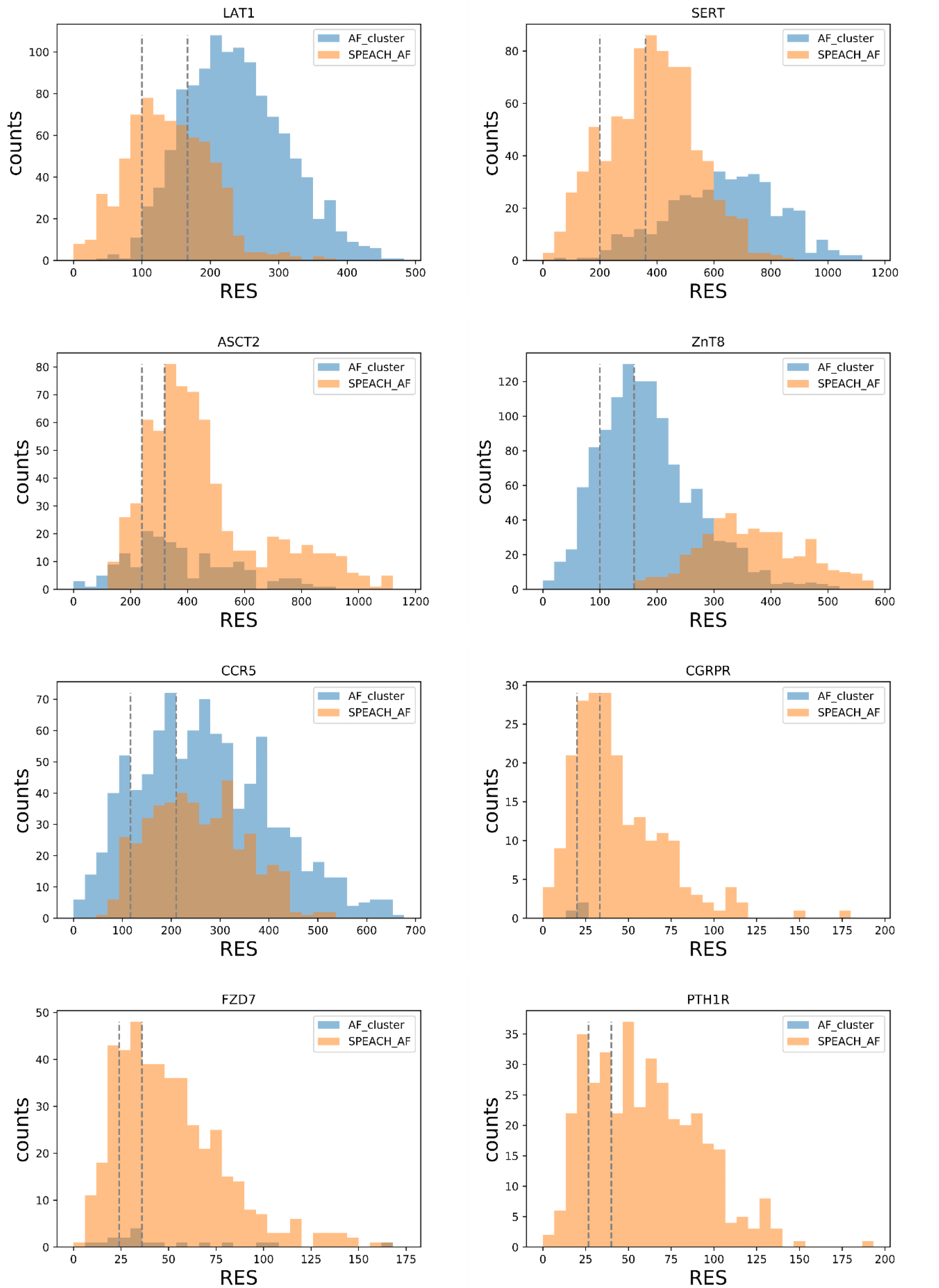
Histograms of RES for AF_cluster and SPEACH_AF. The gray dashed lines demarcate the ∼10% and ∼30% lowest scoring models from both methods combined.

**Figure S15.**
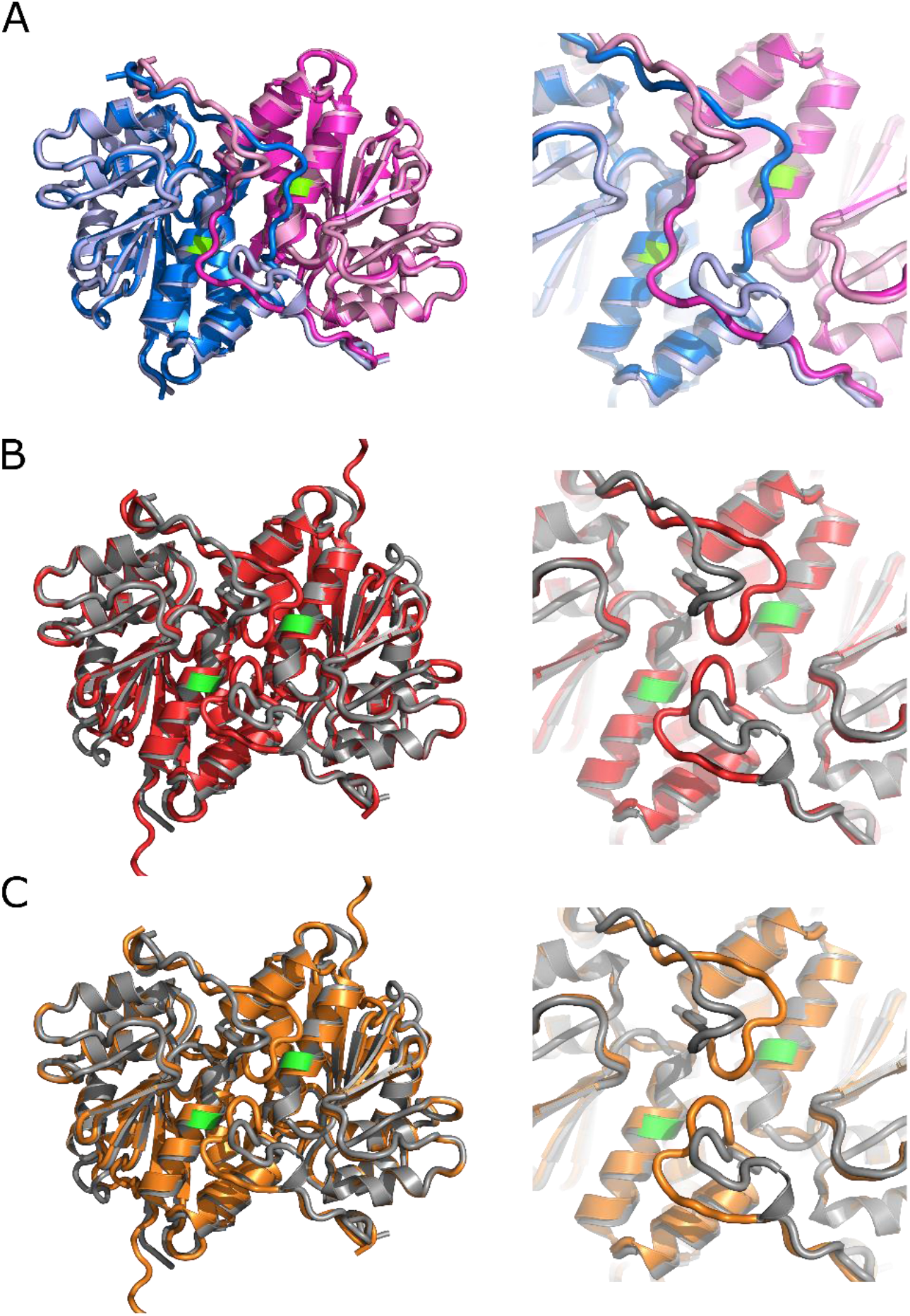
Effect of mutation on isocyanide hydratase. A) Dimer models from CASP 15. The wild-type protein (T1110) is colored slate and magenta. The mutant protein (T1109) is colored lightblue and lightpink. The D183A mutation is colored green. On the right is a close up of the N-terminal tail that switches structure upon mutation. The RMSD for these two structures is 11.3. B) Mutant protein (T1109, grey) and the AlphaFold2 model (red) generated with the mutation across the whole MSA. While different from the crystal structure the AlphaFold2 model has a similar configuration of the N-terminal region. The RMSD between the model and structure is 1.88. C) Mutant protein (T1109, grey) and the AlphaFold2 model (orange) generated with the mutation in the input sequence only. While different from the crystal structure the AlphaFold2 model has a similar configuration of the N-terminal region. The RMSD between the model and structure is 1.87.

**Figure S16.**
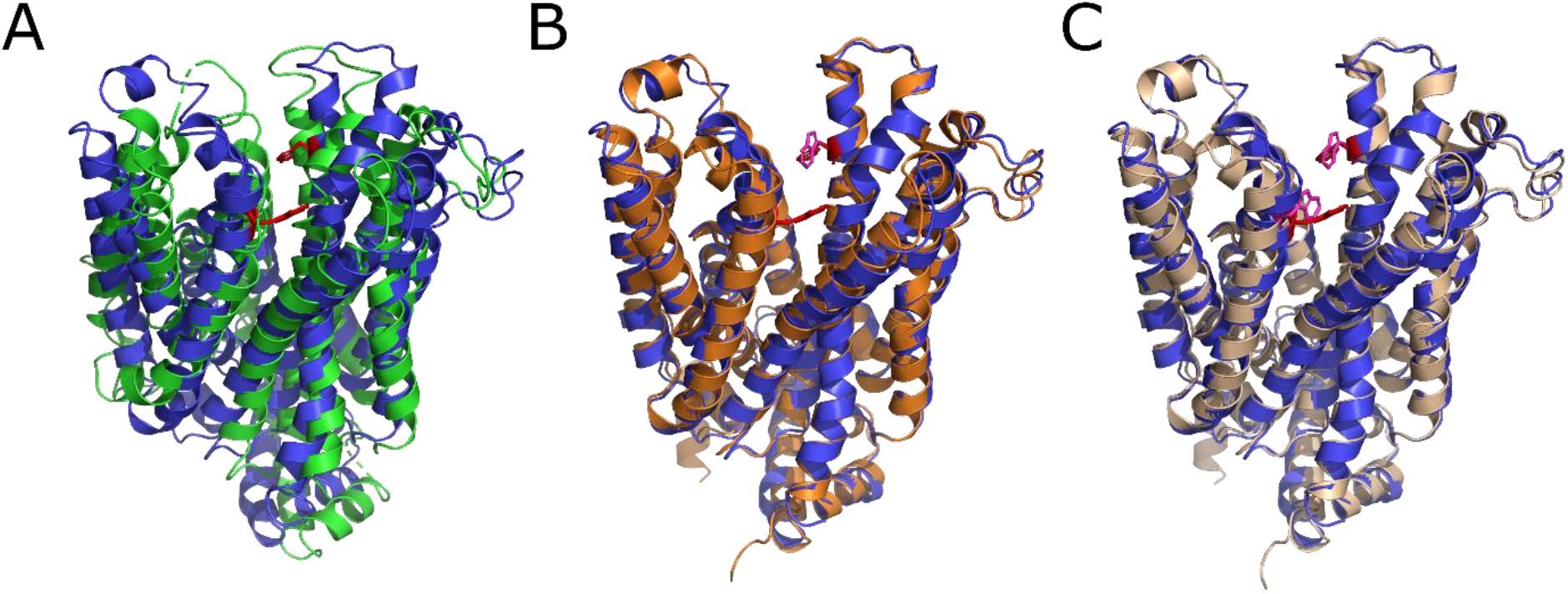
XylE conformational switch. A) Crystal structures of outward-facing (blue, 6n3i) and inward-facing (green, 4ja4) XylE. In red are the mutant residues, G58W/L315W, in 6n3i. The RMSD for these two structures is 4.52. B) Outward-facing crystal structure 6n3i (blue) and outward-facing AlphaFold2 model (orange, best TM-score) with the single G58W mutation (magenta). The RMSD between the model and structure is 1.30. C) Outward-facing crystal structure 6n3i (blue) and outward-facing AlphaFold2 model (wheat, best TM-score) with the G58W/L215W mutation (magenta). The RMSD between the model and structure is 1.31.

**Figure S17.**
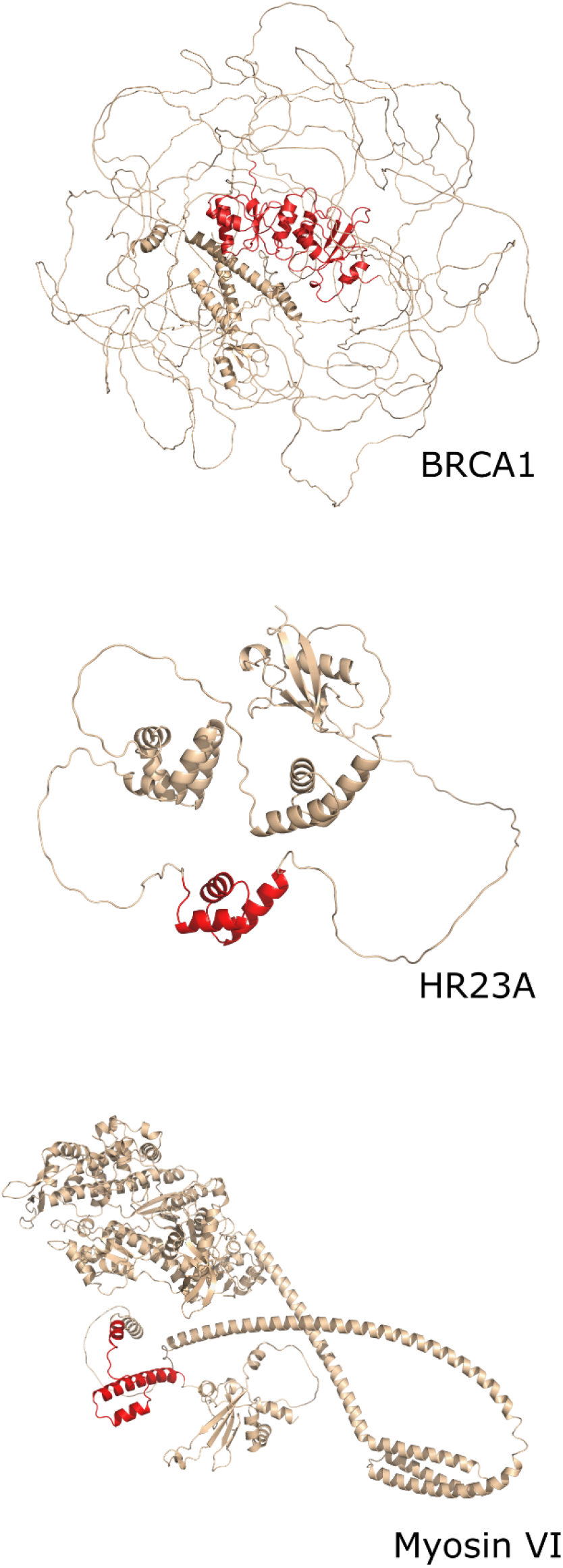
Full length models of BRCA1, HR23A, and Myosin VI. Highlighted in red are the BRCT of BRCA1, UBA of HR23A, and MyUb of Myosin VI domains. The regions in red were used in the comparison to the models of wild-type and mutant domains.

**Figure S18.**
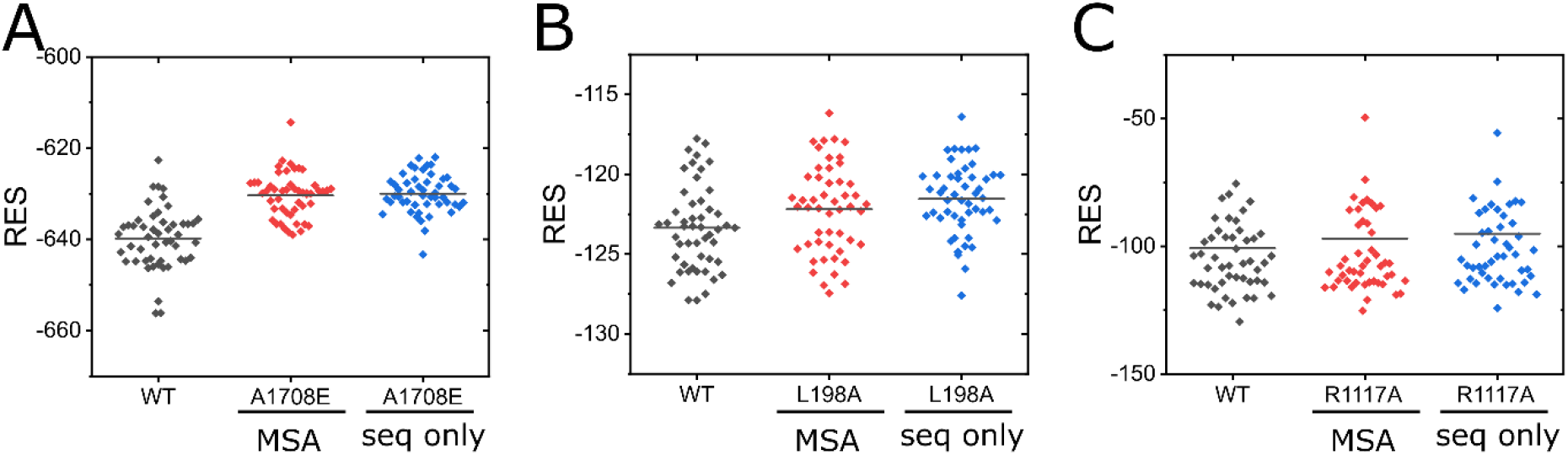
Rosetta Energy Scores for the AlphaFold2 models. For each of the sets of models a t-test was done to ascertain the confidence that the mutant model set is different from the wild-type model set. A) BRCT domain of BRCA1. The p-values for the t-test and Mann-Whitney U test are 2.97E-12 and 1.18E-10, respectively for in the whole MSA and 1.57E-13 and 4.36E-12, respectively for in the input sequence only. B) UBA domain of HR23A. The p-values for the t-test and Mann-Whitney U test are 0.035 and 0.038, respectively for in the whole MSA and 3.63E-4 and 2.55E-4, respectively for in the input sequence only. C) MyUB domain of Myosin VI. The p-values for the t-test and Mann-Whitney U test are 0.531 and 0.797, respectively for in the whole MSA and 0.173 and 0.266, respectively for in the input sequence only.

**Figure S19.**
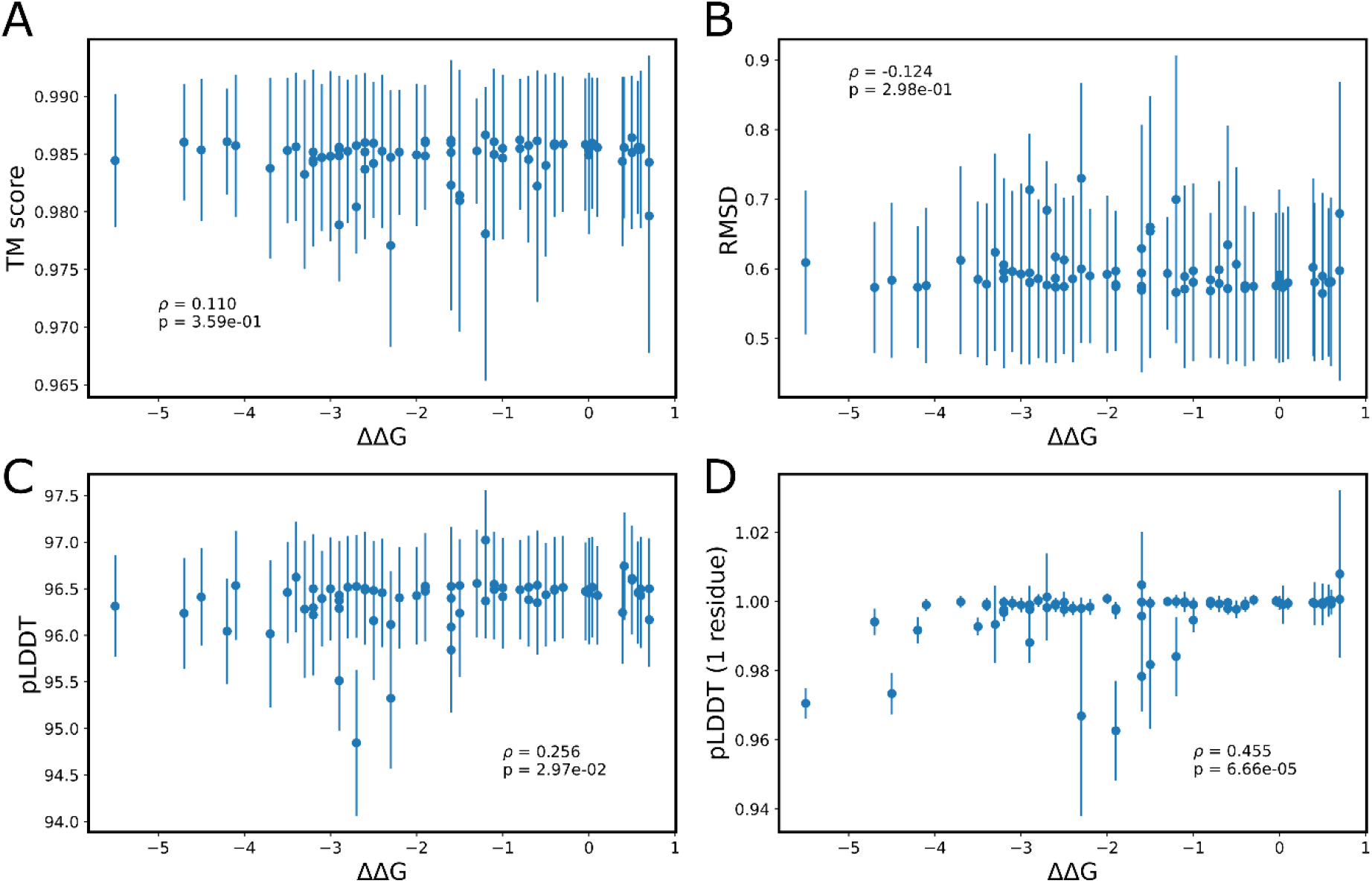
Additional metrics for T4 Lysozyme compared to ΔΔG. A) TM-score and B) RMSD exhibit little to no correlation with ΔΔG and with large p-values suggesting no statistical confidence in the correlation. C) Whole protein pLDDT values exhibits a weak correlation with ΔΔG, though the correlation is not predictive for the effect of almost all of the mutations examined. D) pLDDT value for the mutated residue normalized to the wild-type residue pLDDT value exhibits a significant Pearson’s correlation coefficient, but almost all of the values are at or near 1, making this metric unusable for individual mutations.

**Figure S20.**
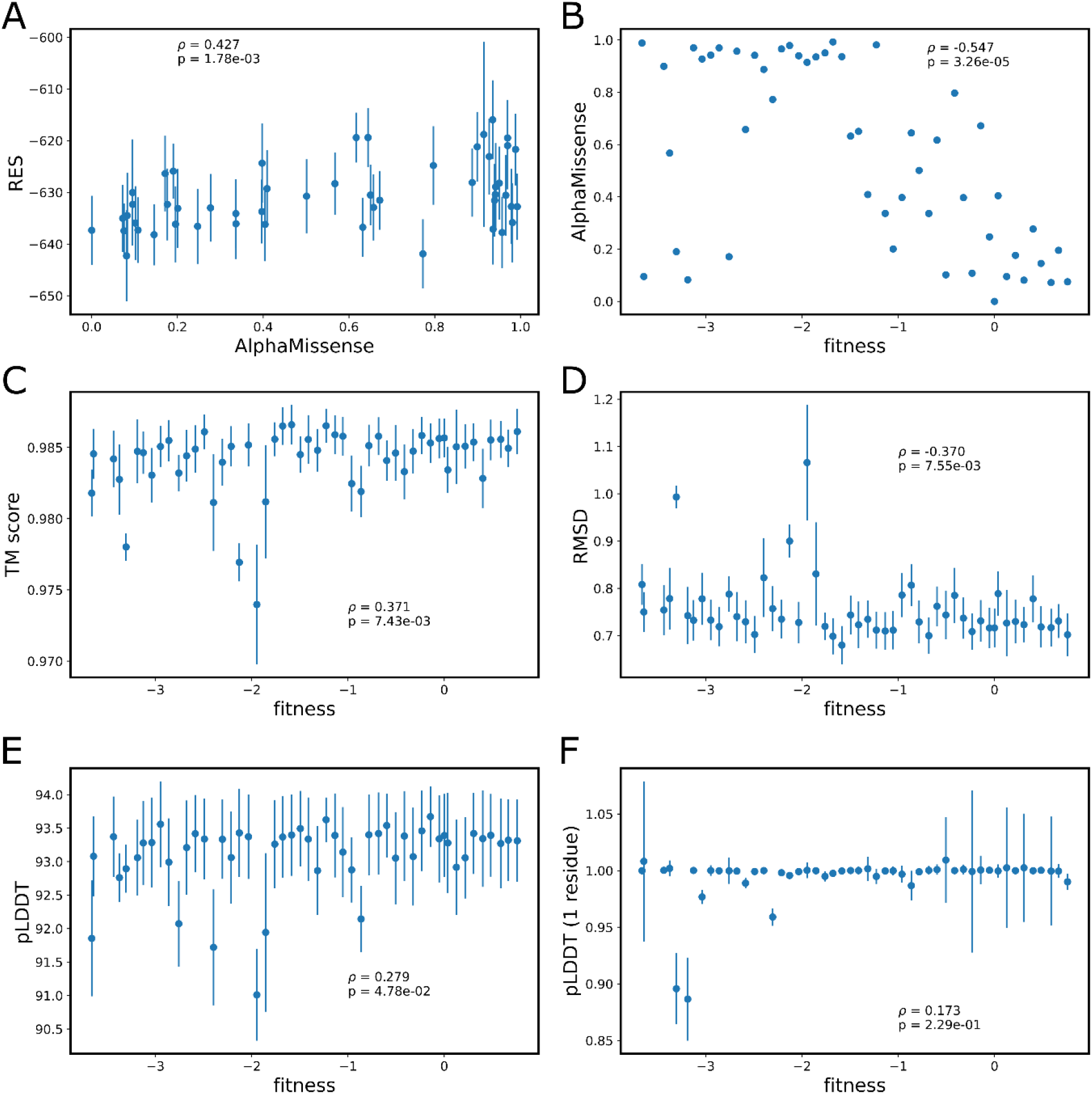
Additional metrics for BRCA1 mutations. A) The RES is moderately correlative to the AlphaMissense score. B) The AlphaMissense score is strongly correlative to fitness. C) The TM score is moderately correlative to fitness. D) The RMSD is moderately correlative to fitness. E) Whole protein pLDDT values exhibits a weak correlation with fitness, though the correlation is not predictive for the effect of almost all of the mutations examined. F) There is no correlation between the pLDDT value for the mutated residue normalized to the wild-type residue pLDDT value relative to the fitness score.

**Table S1:** Rosetta Command Line and Script.

| Table S1: Rosetta Command Line and Script |
| --- |
| Command line:<br>`~/rosetta_bin_linux_2021.16.61629_bundle/main/source/bin/rosetta_scripts.static.linuxgccrelease -s model.pdb -parser:protocol protocol.txt -in:file:spanfile model_span.txt -constrain_relax_to_start_coords -optimization::default_max_cycles 200 -out:path:pdb .` |
| Protocol.txt:<br>`<ROSETTASCRIPTS> <SCOREFXNS> <ScoreFunction name="memb_sc" weights="membrane_highres_Menv_smooth"> <Reweight scoretype="cart_bonded" weight="0.5"/> <Reweight scoretype="pro_close" weight="0"/> </ScoreFunction> </SCOREFXNS> <RESIDUE_SELECTORS> </RESIDUE_SELECTORS> <TASKOPERATIONS> </TASKOPERATIONS> <SIMPLE_METRICS> </SIMPLE_METRICS> <FILTERS> </FILTERS> <MOVERS> <MutateResidue name="res1" target="22A" new_res="SER"/> <FastRelax name="fast_relax" scorefxn="memb_sc" cartesian="True"/> </MOVERS> <PROTOCOLS> <Add mover_name="res1"/> <Add mover_name="fast_relax"/> </PROTOCOLS> <OUTPUT scorefxn="memb_sc"/> </ROSETTASCRIPTS>` |
| Example command line parameters and scripts for Rosetta runs. Command line for running a transmembrane protein initially modelled with SPEACH_AF. The protocol.txt file includes mutating residues back to the wild-type residue prior to minimization and scoring. A MutateResidue line was included for each residue that had been mutated to Alanine for modeling. For soluble proteins the default ScoreFunction was used and the in:file:spanfile was omitted. For examining the effect of single point mutations, the MutateResidue method was omitted. The scoring function for soluble proteins was the default scoring function. |

**Table S2:** T4 Lysozyme Mutants.

| mutant | $\Delta\Delta G$<br>(kcal/mol) | mutant | $\Delta\Delta G$<br>(kcal/mol) | mutant | $\Delta\Delta G$<br>(kcal/mol) | mutant | $\Delta\Delta G$<br>(kcal/mol) |
| --- | --- | --- | --- | --- | --- | --- | --- |
| R96P<sup>1</sup> | -5.5 | R96T<sup>1</sup> | -2.8 | V103A<sup>2</sup> | -1.6 | L99M<sup>3</sup> | -0.4 |
| R96Y<sup>1</sup> | -4.7 | F104A<sup>2</sup> | -2.7 | I100M<sup>3</sup> | -1.6 | L133M<sup>3</sup> | -0.4 |
| R96W<sup>1</sup> | -4.5 | R96M<sup>1</sup> | -2.7 | V87A<sup>2</sup> | -1.5 | R96Q<sup>1</sup> | -0.3 |
| R96F<sup>1</sup> | -4.2 | L91A<sup>2</sup> | -2.6 | I78M<sup>3</sup> | -1.5 | R119E<sup>4</sup> | -0.04 |
| L99A<sup>2</sup> | -4.1 | R96S<sup>1</sup> | -2.6 | A129L<sup>5</sup> | -1.3 | WT | 0 |
| L84A<sup>2</sup> | -3.7 | R96G<sup>1</sup> | -2.6 | I78A<sup>2</sup> | -1.2 | R96K<sup>1</sup> | 0 |
| R96D<sup>1</sup> | -3.5 | I100A<sup>2</sup> | -2.5 | V103M<sup>3</sup> | -1.2 | T115E<sup>6</sup> | 0.04 |
| F153A<sup>2</sup> | -3.4 | R96E<sup>1</sup> | -2.5 | R154E<sup>4</sup> | -1.1 | Q123E<sup>6</sup> | 0.1 |
| L66A<sup>2</sup> | -3.3 | R96A<sup>1</sup> | -2.4 | S90H<sup>6</sup> | -1.1 | V131A<sup>7</sup> | 0.39 |
| L118A<sup>2</sup> | -3.2 | L7A<sup>2</sup> | -2.3 | V111A<sup>2</sup> | -1 | N144D<sup>7</sup> | 0.41 |
| V149A<sup>2</sup> | -3.2 | I17A<sup>2</sup> | -2.3 | K135E<sup>4</sup> | -1 | K16E<sup>4</sup> | 0.5 |
| R96L<sup>1</sup> | -3.2 | L121A<sup>2</sup> | -2.2 | L91M<sup>3</sup> | -0.8 | N144E<sup>6</sup> | 0.5 |
| R96H<sup>1</sup> | -3.1 | R96V<sup>1</sup> | -2 | L121M<sup>3</sup> | -0.8 | A82P<sup>7</sup> | 0.57 |
| R96N<sup>1</sup> | -3 | A129M<sup>5</sup> | -1.9 | K147E<sup>4</sup> | -0.7 | S38D<sup>7</sup> | 0.6 |
| L33A<sup>2</sup> | -2.9 | M106A<sup>2</sup> | -1.9 | L118M<sup>3</sup> | -0.7 | N116D<sup>7</sup> | 0.6 |
| M102A<sup>2</sup> | -2.9 | L84M<sup>3</sup> | -1.9 | K85A<sup>1</sup> | -0.6 | E108V<sup>8</sup> | 0.7 |
| R96C<sup>1</sup> | -2.9 | M6A<sup>2</sup> | -1.6 | F153M<sup>3</sup> | -0.6 | I3L<sup>7</sup> | 0.7 |
| R96I<sup>1</sup> | -2.9 | I50A<sup>2</sup> | -1.6 | D89A<sup>1</sup> | -0.5 |  |  |

**Table S3:** BRCA1 Mutants in the BRCT Domain.

| Table S3: BRCA1 Mutants in the BRCT Domain |  |  |  |  |  |  |  |  |
| --- | --- | --- | --- | --- | --- | --- | --- | --- |
| mutant | fitness<sup>a</sup> | AM<sup>b</sup> | mutant | fitness<sup>a</sup> | AM<sup>b</sup> | mutant | fitness<sup>a</sup> | AM<sup>b</sup> |
| G1738R | -3.66162 | 0.9885 | R1753S | -2.12872 | 0.9788 | Q1779P | -0.5975 | 0.6168 |
| A1823S | -3.64283 | 0.0953 | F1734S | -2.03381 | 0.9397 | T1816K | -0.5051 | 0.1017 |
| F1761I | -3.43496 | 0.8991 | Q1811P | -1.9469 | 0.9144 | G1710V | -0.41422 | 0.7969 |
| V1810G | -3.37388 | 0.5673 | T1691K | -1.85493 | 0.9352 | V1792L | -0.32322 | 0.3969 |
| A1823P | -3.30465 | 0.1901 | F1668S | -1.76069 | 0.9506 | V1646L | -0.23303 | 0.1078 |
| A1823T | -3.18652 | 0.0828 | W1718R | -1.67812 | 0.9925 | N1774I | -0.14358 | 0.6718 |
| A1752P | -3.12678 | 0.97 | H1686Q | -1.58542 | 0.9361 | M1663I | -0.05254 | 0.247 |
| A1752E | -3.03783 | 0.9271 | V1842A | -1.49481 | 0.6322 | WT | 0 | 0 |
| I1766N | -2.94743 | 0.9422 | G1788A | -1.41213 | 0.6502 | K1759N | 0.03674 | 0.4042 |
| R1751P | -2.86079 | 0.9699 | D1813Y | -1.3163 | 0.4088 | E1817K | 0.126227 | 0.0952 |
| P1812A | -2.75731 | 0.171 | C1768R | -1.22753 | 0.9812 | R1744K | 0.217324 | 0.1763 |
| P1749Q | -2.67599 | 0.9571 | S1655A | -1.13543 | 0.3365 | T1802P | 0.305815 | 0.0812 |
| A1752T | -2.58579 | 0.6574 | D1778Y | -1.05268 | 0.2006 | E1781A | 0.400263 | 0.277 |
| L1839S | -2.49144 | 0.9413 | C1767G | -0.96147 | 0.3974 | T1658S | 0.483314 | 0.1454 |
| C1697F | -2.39626 | 0.8875 | I1855K | -0.86384 | 0.6445 | Q1756E | 0.587609 | 0.0721 |
| H1746Y | -2.30475 | 0.7724 | V1741L | -0.78295 | 0.5011 | H1672L | 0.664879 | 0.1954 |
| Y1703N | -2.21296 | 0.9658 | T1677P | -0.68274 | 0.3362 | M1650V | 0.758385 | 0.0752 |
| a: fitness values from ProteinGym (https://proteingym.org/) based on data from Findlay GM, Daza RM, Martin B, Zhang MD, Leith AP, Gasperini M, Janizek JD, Huang X, Starita LM, Shendure J (2018) Accurate classification of BRCA1 variants with saturation genome editing. Nature 562:217-222.<br>b: AlphaMissense values from Cheng J, Novati G, Pan J, Bycroft C, Žemgulytė A, Applebaum T, Pritzel A, Wong LH, Zielinski M, Sargeant T, Schneider RG, Senior AW, Jumper J, Hassabis D, Kohli P, Avsec Ž. Accurate proteome-wide missense variant effect prediction with AlphaMissense. Science 381:eadg7492. |  |  |  |  |  |  |  |  |

**Table S4:** Breakdown of the Initial Screen of AlphaFold2-generated Models.

| Table S4: Breakdown of the Initial Screen of AlphaFold2-generated Models<sup>a</sup> |  |  |  |  |
| --- | --- | --- | --- | --- |
| protein | AFC_3<sup>b</sup> | AFC_7<sup>b</sup> | AFC_11<sup>b</sup> | SPEACH_AF |
| AK | 210/82 | 558/13 | 1109/8 | 45/45 |
| RBP | 1710/0 | 325/0 | 95/0 | 2/0 |
| MCT1 | 1191/0 | 162/3 | 66/2 | 2/0 |
| STP10 | 745/4 | 27/0 | 4/0 | 0/0 |
| MurJ | 736/18 | 84/0 | 46/0 | 170/168 |
| PfMATE | 77/5 | 0/0 | 0/0 | 2/0 |
| Lat1 | 1294/0 | 241/0 | 56/3 | 0/0 |
| SERT | 1324/4 | 498/2 | 266/4 | 0/0 |
| ASCT2 | 1185/0 | 258/0 | 64/0 | 3/0 |
| ZnT8 | 1301/0 | 139/0 | 20/0 | 15/15 |
| CCR5 | 665/74 | 132/53 | 49/23 | 5/3 |
| CGRPR | 520/0 | 205/0 | 137/0 | 173/149 |
| FZD7 | 550/0 | 163/0 | 100/0 | 2/2 |
| PTH1R | 105/0 | 50/0 | 35/0 | 63/21 |
| a: Shown are the number of models that failed the initial two-step screen and the number of those models that are misfolded/misthreaded. |  |  |  |  |
| b: AFC_3, AFC_7, and AFC_11 utilizing AF_cluster with the minimum number of sequences set to 3, 7, and 11 respectively. |  |  |  |  |

